# Adaptation in cone photoreceptors contributes to an unexpected insensitivity of On parasol retinal ganglion cells to spatial structure in natural images

**DOI:** 10.1101/2021.06.29.450295

**Authors:** Zhou Yu, Maxwell H. Turner, Jacob Baudin, Fred Rieke

## Abstract

Neural circuits are constructed from nonlinear building blocks, and not surprisingly overall circuit behavior is often strongly nonlinear. But neural circuits can also behave near linearly, and some circuits shift from linear to nonlinear behavior depending on stimulus conditions. Such control of the linearity or nonlinearity of circuit behavior is fundamental to neural computation. Here we study a surprising stimulus dependence of the responses of On (but not Off) parasol retinal ganglion cells: these cells respond nonlinearly to spatial structure in temporally-modulated grating stimuli but linearly to spatial structure in flashed gratings and natural visual inputs. We show that this unexpected response linearity can be explained by a shift in the balance of excitatory and inhibitory synaptic inputs that originates at least in part from adaptation in the cone photoreceptors. More generally, this highlights how subtle asymmetries in signaling - here in the cone signals - can qualitatively alter circuit computation.

## Introduction

Components of neural circuits often transform neural signals nonlinearly. Common nonlinear relations include those between a sensory stimulus and the response of a primary sensory receptor (Hudspeth and Corey, 1977; Baylor et al., 1987), between presynaptic voltage and synaptic release (Katz and Miledi, 1967; Huang and Neher, 1996), and between membrane potential and action potential generation (Hodgkin and Huxley, 1952). Neural computation relies on the judicious control of these nonlinearities - in some cases to make overall circuit behavior near linear despite sharp deviations from linearity in the underlying components (Werblin, 2010). In some instances, linear circuit behavior emerges because signals are small and the underlying circuit mechanisms are not modulated sufficiently strongly to reveal their nonlinearities. In others, multiple nonlinear mechanisms act cooperatively to produce linear circuit behavior. Understanding how such control is exerted and when a neural circuit operates near linearly has important consequences for constructing models of the nervous system and can provide a strong constraint for what kind of processing a circuit performs.

Retinal output neurons (i.e. retinal ganglion cells or RGCs) have long been classified by whether or not they linearly integrate signals across space (reviewed by Field and Chichilnisky, 2007; Sanes and Masland, 2015). Although this binary classification of spatial sensitivity is oversimplified, it has proven quite useful. The responses of spatially-linear RGCs are proportional to the total light incident on their receptive field, whereas spatially-nonlinear RGCs are also sensitive to the spatial distribution of light within the receptive field (Enroth-Cugell and Robson, 1966; Hochstein and Shapley, 1976). Classical tests of spatial integration used temporally modulated grating stimuli. These stimuli allow a clear prediction to be made for a linear RGC: a spatially-linear RGC should produce no response because the light and dark bars that form the grating are equal and opposite in contrast and hence should cancel upon summation over space. RGCs that respond to gratings exhibit nonlinear spatial integration; such spatial nonlinearity can be explained if a nonlinearity at the output synapses of the bipolar cells presynaptic to a RGC causes responses to the light and dark bars not to cancel (Demb et al., 1999) (Figure 1A).

**Figure 1:**
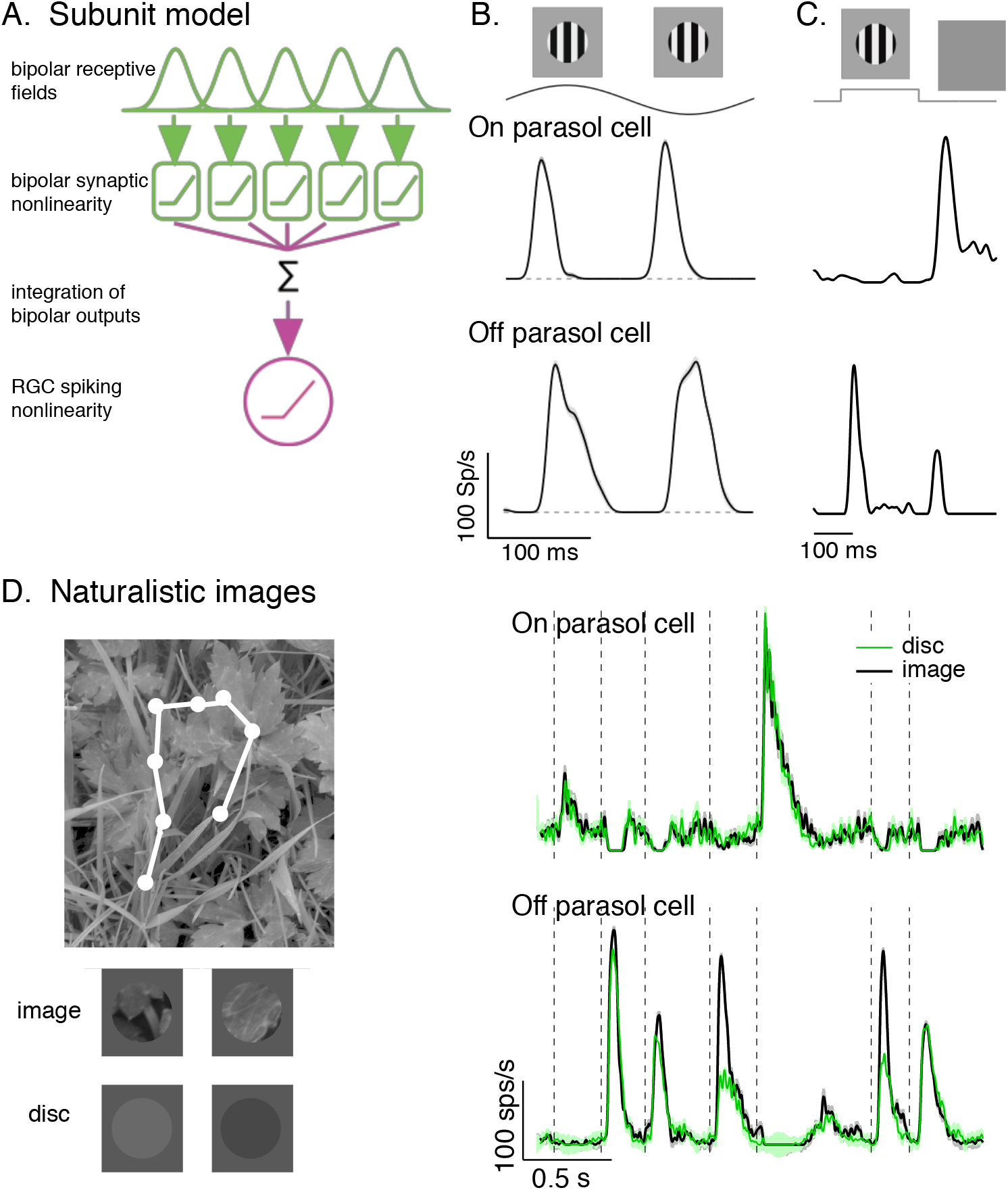
Differences in spatial integration for gratings and naturalistic stimuli. A. Standard subunit model often used to account for nonlinear spatial integration by RGCs. B. Both On and Off parasol cells show strong, nonlinear responses to contrast reversing gratings. C. On and Off parasol cell responses to flashed gratings differ. D. On but not Off parasol cells show near-linear responses to natural movies. Top left shows image with eye movement trajectory in white. Bottom left shows two natural image movie frames and corresponding linear-equivalent discs. (Right) Black trace shows responses to a natural movie generated using the DOVES database (Van Der Linde et al., 2009). Movies were restricted to the receptive field center with a circular aperture. Green shows responses to a linear-equivalent movie, in which spatial structure within the receptive field center was replaced with a uniform disc with an intensity equal to the weighted average intensity within the receptive field center. The weighting was determined by a gaussian fit to the dependence of the response on the size of a test spot (see Turner and Rieke, 2016 for details).

In a simple model like that in Figure 1A, the bipolar cell synaptic nonlinearities that shape responses to temporally-modulated gratings also explain sensitivity to spatial structure in other stimuli. However, many stimuli encountered by the retina - e.g. natural images coupled with eye movements - contain more complex spatial and temporal structure than periodic spatial gratings. Further, retinal circuits contain nonlinear circuit elements in addition to bipolar cell synapses. These two observations suggest that insights about spatial integration from grating stimuli may not generalize well to other stimuli, particularly natural ones. Indeed, we found previously that On parasol (magnocellular-projecting) RGCs exhibited strongly nonlinear spatial integration for temporally-modulated grating stimuli but near-linear spatial integration for natural inputs (Turner and Rieke, 2016). Off parasol cells exhibited nonlinear spatial integration for both stimulus classes. Here, we explore the mechanistic basis of this On/Off asymmetry and its consequences for natural scene encoding. We find that: (1) flashed gratings and natural inputs recruit large inhibitory synaptic inputs to On parasol RGCs that cancel spatial nonlinearities in excitatory synaptic inputs and suppress sensitivity of the spike output to spatial structure. (2) Adaptation in the cone photoreceptors creates subtle asymmetries in the circuits controlling On parasol excitatory and inhibitory synaptic inputs. These asymmetries, magnified by circuit nonlinearities, can largely account for the differences in spatial integration between temporally-modulated gratings and flashed gratings or natural inputs. And, (3) differences in spatial sensitivity of On and Off cells may enhance population coding of specific image features at the expense of others.

## Results

We begin with evidence that the insensitivity of responses of On parasol RGCs to spatial structure in natural inputs originates in the integration of excitatory and inhibitory synaptic inputs. We then show that adaptation in the cone photoreceptors plays a central role in controlling the balance of excitatory and inhibitory inputs, and that differences in On parasol responses to gratings and natural inputs can be largely explained by differences in the way that cones adapt to these stimuli. Finally, we explore how the difference in spatial integration between On and Off parasol RGCs impacts the encoding of specific stimulus properties.

### Unexpected linearity of On parasol cell responses to spatial structure in natural images

A classic test for nonlinear spatial integration is to periodically modulate a spatial grating restricted to the RGC receptive field center and measure the average response to a single modulation cycle. Nonlinear spatial integration causes a response at twice the temporal frequency of modulation -- i.e. a “frequency-doubled” or F2 response (Enroth-Cugell and Robson, 1966; Hochstein and Shapley, 1976). As observed previously, cycle-averaged responses of both On and Off parasol RGCs showed strong frequency-doubled responses to contrast-reversing gratings (Petrusca et al., 2007; Crook et al., 2008; Cafaro and Rieke, 2013) (Figure 1B). These responses are consistent with a nonlinearity in the bipolar output signals which creates subunits in the On parasol receptive field (Figure 1A).

The model in Figure 1A, however, failed to predict On parasol responses to spatial structure in several other stimuli. First, flashed gratings produced very different responses in On and Off parasol cells (Figure 1C). Specifically, Off parasol cells, as predicted from the model in Figure 1A, responded at both the onset and offset of a flashed grating. On parasol cells, however, responded minimally at the grating onset and strongly at the offset.

Second, On parasol cells responded weakly to spatial structure in natural inputs (Figure 1D). As in previous work (Turner and Rieke, 2016; Turner et al., 2018), we probed spatial integration of natural images by comparing responses to spatially-structured natural inputs with responses to spatially-uniform stimuli with temporal variations chosen to match those of the natural inputs (see below). We designed these stimuli to explore responses to naturalistic spatial and temporal structure in the receptive field center; there are clearly other aspects of real visual inputs that they do not capture.

Natural movie stimuli mimicked the retinal input during free viewing by translating a natural image across the retina according to measured human eye movements (Van Der Linde et al., 2009) (Figure 1D, top). The intensity of each frame of the linear-equivalent disc movie was calculated by linearly integrating the intensity of the corresponding frame of the natural movie weighted by an estimate of the cell’s linear receptive field center. By construction, a cell that integrates inputs linearly over the receptive field center would produce identical responses to the original image and the disc. Both naturalistic and linear-equivalent stimuli were restricted to the receptive field center using an aperture, as determined for each recorded cell from the dependence of response amplitude on spot size (Turner and Rieke, 2016). Figure 1D shows example images and corresponding linear equivalent discs.

As reported previously, Off parasol RGCs responded differently to natural and linear-equivalent movies (compare green and black traces in Figure 1D, right; Turner and Rieke, 2016). This difference indicates that Off parasol RGCs respond nonlinearly to spatial structure in natural visual inputs, as is expected from their responses to contrast-reversing and flashed gratings. On parasol RGCs, however, showed similar responses to natural and linear-equivalent movies (Figure 1D, right); thus, unlike their responses to temporally-modulated gratings, On parasol RGCs integrated spatial structure in natural inputs within their receptive field center linearly or near-linearly. Spatial integration by On and Off parasol cells differed across a broad range of light levels (Turner and Rieke, 2016). The difference in sensitivity of On parasol cells to spatial structure in temporally-modulated gratings and natural images is unexpected from standard subunit models for spatial integration (such as the model in Figure 1A). It is also interesting given the importance of these cells in visual perception and the possible functional implications of such a strong asymmetry between matched On and Off RGC types.

### On parasol excitatory inputs elicited by natural inputs are more nonlinear than spike outputs

Interpreting responses to natural image movies like those in Figure 1D is complicated by possible history dependence - e.g. the response at a given time could reflect the onset of a new image feature and/or the offset of a previous feature. To simplify the correspondence between image features and responses, we turned to flashed image patches (Figure 2). All patches were flashed starting from the same gray background, and hence differences in response can be attributed to the structure in the patch itself rather than the past history of the stimulus. To focus on patches that elicited spike responses from On parasol RGCs, we selected patches with a higher luminance in the receptive field center compared to the gray background.

**Figure 2:**
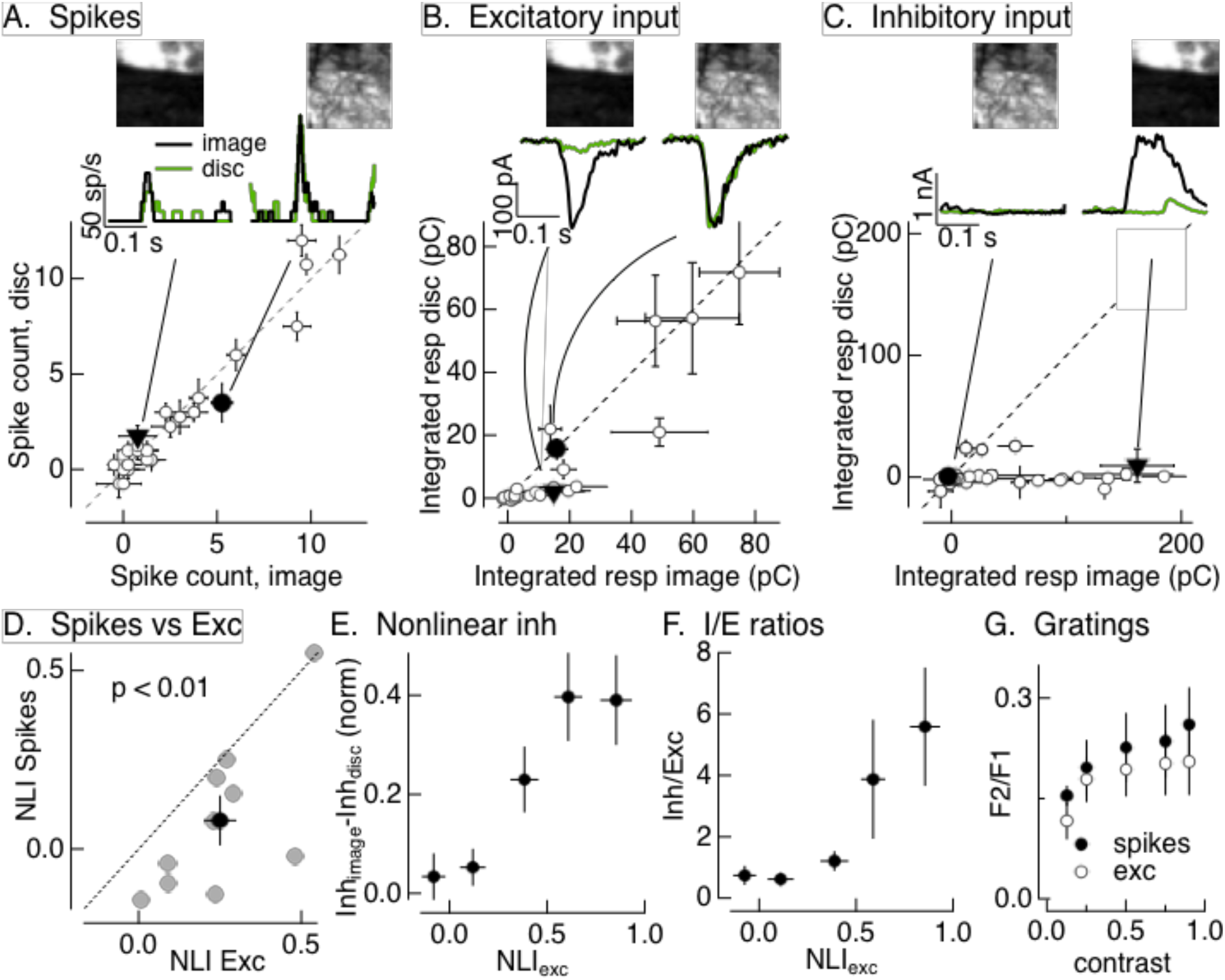
Synaptic integration leads to unexpected linearity of On parasol responses to natural images. A. Spike count for On parasol responses to image patches and corresponding linear equivalent stimuli. Each stimulus was flashed for 250 ms and spikes counted during that time. Peri-stimulus time histograms for the two highlighted points are at the top along with the corresponding images; the black trace is the response to the image patch and the green trace is the response to the linear equivalent stimulus. B. Excitatory synaptic inputs for the same cell and image patches as in A. Responses are the integrated current during the stimulus presentation. The same two image patches are highlighted. C. Inhibitory synaptic input for the same cell and image patches. D. Excitatory inputs are more nonlinear than spike output. Comparison of nonlinearity index (NLI) for spike response with that for excitatory input for 9 On parasol cells. E. Patches that recruit nonlinear excitatory input also recruit nonlinear inhibitory input. Nonlinear inhibitory input (image response - disc response) plotted against excitatory NLI. Results combined from 8 On parasol cells. F. Ratio of inhibitory input to excitatory input increases for patches that elicit strong nonlinear excitatory input. Combined results from 8 On parasol cells. G. On parasol spikes and excitatory inputs exhibit similar nonlinear spatial integration in response to contrast-reversing gratings across a range of contrasts. The strength of nonlinear spatial integration was summarized as the ratio of the frequency doubled or F2 response measured for a split-field grating to the F1 response measured for a modulated spot.

For each cell, we recorded spike responses, excitatory synaptic inputs and inhibitory synaptic inputs (Figure 2A-C; see Methods) elicited by a set of image patches and corresponding linear-equivalent discs. Responses were quantified by integrating the response over the time that the stimulus was presented. Consistent with the responses to time-varying naturalistic stimuli in Figure 1D, On parasol RGCs generated similar spike responses to flashed natural images and linear-equivalent discs (Figure 2A).

Two aspects of the responses to flashed image patches suggested that inhibitory synaptic input contributes to the linearity of On parasol RGC spike responses. First, two image patches that elicited similar amplitude excitatory input could elicit quite different spike responses; e.g., the black traces at the top of Figure 2A and B are from the same two image patches. The excitatory inputs for these patches were similar in amplitude, but the spike responses differed considerably. This is consistent with inhibitory input suppressing the response to the patch on the left relative to that on the right. Second, excitatory inputs exhibited clear nonlinear spatial integration, as indicated by comparing responses to flashed image patches and corresponding linear equivalent discs (black and green responses at the top of Figure 2B). Excitatory inputs to many image patches exceeded responses to linear equivalent discs, e.g. the cluster of points below the unity line in Figure 2B. Spike responses of the same cell to the same image patches are much more linear, as indicated by the similarity of the black and green traces at the top of Figure 2A and the similarity of the image and disc responses across patches illustrated in the bottom panel.

To test the generality of the difference in spatial integration between excitatory inputs and spike output across cells and images, we summarized responses with a nonlinearity index. This index measures the normalized difference between responses to the image and disc

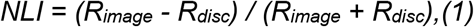

where *R_image_* is the response to the image and *R_disc_* the response to the disc. NLIs varied considerably across different cells and images (Figure 2D), with some images systematically producing positive NLIs for both spike output and excitatory input. Across cells, however, the nonlinearity index (NLI) for excitatory inputs was significantly larger than that for spike output (Figure 2D; 0.25 ± 0.05 vs 0.08 ± 0.07, mean ± SEM, p < 0.01, n=10). This is not due to large NLIs in a small subset of image patches; instead, more than half of the patches from a typical image showed nonlinear spatial integration for excitatory inputs and near-linear spatial integration for spike outputs (as in Figure 2A,B).

We could not use the NLI to test for a similar difference in nonlinear spatial integration for contrast-reversing gratings since the linear equivalent disc in this case would have no contrast and hence elicit no response. Instead, we compared the response to a split-field grating with that to a spatially-uniform spot in the receptive field center modulated at the same frequency and contrast as the grating. We compared the nonlinear (frequency-doubled or F2) response to the grating with the linear response (the response at the modulation frequency or F1 response) to the spot. The F2/F1 ratio was similar for spikes and excitatory inputs (Figure 2G; see also Turner and Rieke, 2016) across a range of contrasts. Thus, for sinusoidally-modulated gratings, nonlinear spatial integration in the On parasol spike output closely follows that in the excitatory inputs. This differs markedly from the situation for both flashed (Figure 2D) and time-varying (Figure 1D) natural images.

### Natural stimuli recruit strong inhibitory input to ganglion cells

A logical hypothesis that follows from the results above is that natural images recruit strong inhibitory synaptic input to On parasol cells that cancels nonlinear excitatory input. The results described below support this hypothesis.

Figure 2C shows inhibitory inputs in response to the same image patches in the same cell as the spikes and excitatory inputs in Figure 2A,B. Inhibitory synaptic input exhibited strongly nonlinear spatial integration, as indicated by the points that fall below the unity line in Figure 2C. Image patches that elicit similar excitatory input can elicit very different inhibitory input; e.g., the filled points and corresponding black traces in the top panels of Figure 2B,C. Further, this difference corresponds to the difference in spike response (black traces in Figure 2A; note that the left/right positions of the matching panels at the top of Figures 2A and C differ). Patches that elicit highly nonlinear excitatory input can also elicit highly nonlinear inhibitory input (e.g. the filled triangles in Figure 2B,C) -- i.e. spatial structure in natural image patches recruits both excitatory and inhibitory input. These observations are consistent with cancellation of nonlinear excitatory input by inhibitory input and the insensitivity of the spike response to spatial structure in patches that elicit strongly nonlinear excitatory input (e.g. the filled triangle and corresponding responses in Figure 2A).

To test the generality of these observations, we quantified the relationship between excitatory and inhibitory input across patches and RGCs. To effectively cancel nonlinear excitatory input, inhibitory input should overlap in time with excitatory input and have at least two additional properties. First, nonlinear inhibitory input should be recruited by the same image patches that elicit nonlinear excitatory input. To test this, we compared the NLI for excitatory input for each individual patch as in Equation 1 with the difference between the inhibitory input elicited by the image and disc (we did not use NLIs for inhibitory input because they tended to be either 0 or 1, obscuring any graded dependence on excitatory NLI). Nonlinear inhibitory input increased systematically with increasing excitatory NLI (Figure 2E). Thus, patches that elicited nonlinear excitatory input also elicited nonlinear inhibitory input.

Second, inhibitory input should be sufficiently large to effectively cancel nonlinearities in excitatory input. To test this, we measured the ratio of the amplitude of inhibitory input to excitatory input (the I/E ratio) as a function of the nonlinearity in excitatory input (Figure 2F). Given the ~3-fold larger driving force on excitatory input near the threshold for spike generation, inhibitory inputs (measured as currents at the reversal potential for excitatory inputs) need to be at least 3-fold larger than excitatory inputs to have a substantial impact on spike output. Patches that elicited near-linear excitatory input corresponded to I/E ratios <1; in this case inhibitory input is unlikely to substantially affect spike output. Patches that elicited strongly nonlinear excitatory input, however, corresponded to I/E ratios >3; for these patches inhibitory input is sufficiently large to impact spike output.

Collectively, the results of Figure 2 indicate that (1) excitatory synaptic input to On parasol cells exhibits larger spatial nonlinearities than spike outputs, and (2) inhibitory synaptic input has the required properties to compensate for nonlinearities in excitatory input and account for the linearity of the spike response. We next characterize inhibitory input in more detail and investigate why it does not linearize responses to grating stimuli as it does responses to natural images.

### Crossover inhibitory input linearizes On parasol responses

Inhibitory synaptic input to On parasol RGCs has both feedforward and crossover components (Figure 3A) (Cafaro and Rieke, 2013; Crook et al., 2014). Feedforward inhibitory input, like excitatory input, is generated by increases in light intensity (Figure 3A, top), whereas crossover inhibitory input is generated by decreases in light intensity (Figure 3A, middle). This means that a spatially uniform increase in light intensity increases both excitatory input and feedforward inhibitory input, while a spatially uniform decrease in intensity decreases excitatory input and increases crossover inhibitory input.

**Figure 3:**
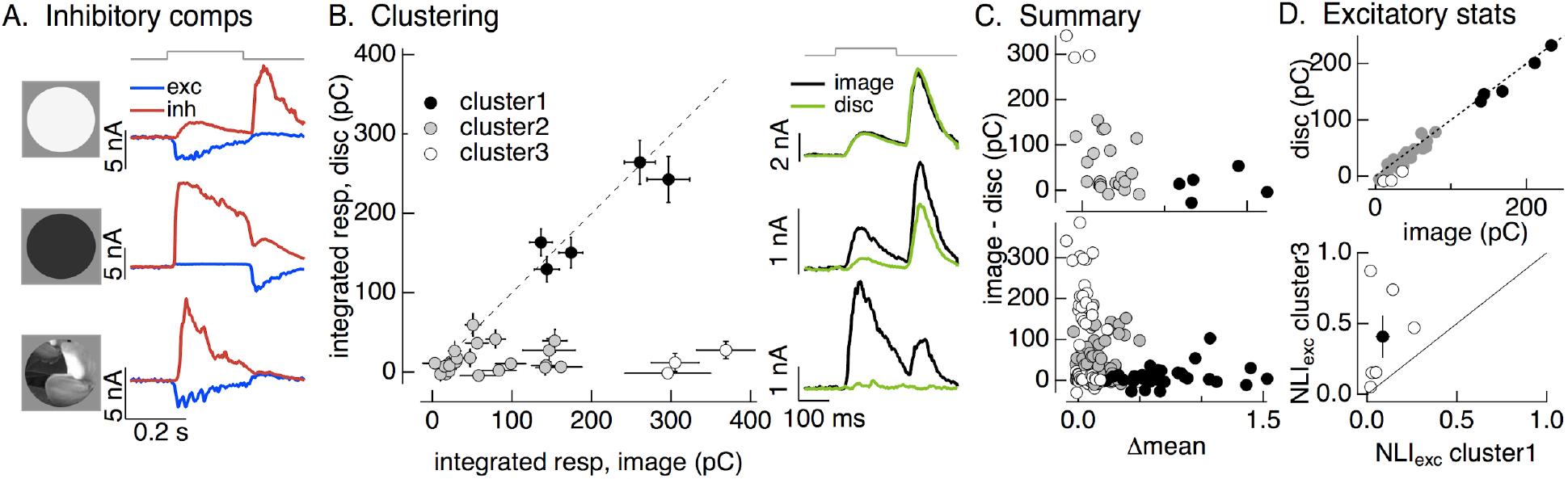
Identifying the components of inhibitory input to On parasol cells elicited by natural image patches. A. Light increments (top) produce an increase in excitatory input (blue) and a small, delayed increase in feedforward inhibitory input (red). Light decrements (middle) produce a decrease in excitatory input and a large increase in crossover inhibitory input. Different regions of spatially-structured inputs (bottom) can produce large crossover inhibitory input that coincide with increases in excitatory input. B. Clustering of inhibitory synaptic inputs elicited by image patches. Clustering based on response time course. Insets at right show average responses to the image (black) and corresponding linear equivalent disc (green) for each cluster. Cluster 1 is dominated by feedforward inhibitory input and shows linear spatial integration. Cluster 3 is dominated by crossover inhibitory input and shows strongly nonlinear spatial integration. C. Summary of relation between nonlinear inhibitory input (image - disc) and mean luminance for each of the clusters from B (top), and summary across cells (bottom). D. (top) Excitatory responses to image patches and corresponding linear equivalent discs for each cluster from B. (bottom) Comparison of nonlinearity indices for excitatory inputs corresponding to clusters 1 and 3.

For spatially-uniform stimuli, feedforward inhibitory input is not sufficiently large to have a sizable impact on spike responses to light increments (I/E ratios are generally less than 1), and crossover inhibitory input has little impact on responses to decrements because the decrease in excitatory input alone is sufficient to eliminate spiking (Cafaro and Rieke, 2013). However, different regions of spatially-structured stimuli can elicit different types of synaptic input - e.g. bright regions of a stimulus can cause an increase in excitatory input while dark regions can cause an increase in crossover inhibitory input (Figure 3A, bottom). This can cause the (potentially large) crossover inhibitory input to overlap in time with excitatory input and hence impact spike output (Cafaro and Rieke, 2013).

To determine which components of inhibitory input were elicited by different natural image patches, we used a principal components-based approach to cluster inhibitory responses (see Methods); clustering used only the time course of the responses to the flashed image patches, and did not use any information about the stimuli or responses to the linear equivalent discs. We set the target number of clusters to three as this was the minimum needed to separate responses with clearly different time courses. The three clusters defined by this approach had consistently different spatial integration properties. Figure 3B (left) compares inhibitory synaptic inputs elicited by flashed image patches and corresponding linear-equivalent discs for each response in each cluster for an example RGC, and Figure 3B (right) shows the time course of average image and disc responses in each cluster. Responses in the first cluster showed linear spatial integration -- i.e. responses to image patches and linear equivalent discs were near identical. Responses in the third cluster exhibited strongly nonlinear spatial integration -- with much larger responses to images than to the corresponding linear equivalent discs. Responses in the second cluster were smaller in amplitude and had more variable spatial integration properties than the others; we omitted these responses from the remaining analysis to focus on responses that were associated with either clearly linear or clearly nonlinear spatial integration.

As a control, we applied the clustering procedure used for images to responses to increment and decrement spots. We retained the cluster definitions determined for the images - i.e. responses to increment and decrement spots were assigned to the clusters defined based on the time course of responses to flashed image patches. High contrast increment spots, which elicit feedforward inhibitory input, led to responses in the first cluster, intermediate contrast increments led to responses in the second cluster, and decrement spots, which elicit crossover inhibitory input, led to responses in the third cluster (Figure 3 - Figure Supplement 1). Hence the clustering approach separates responses associated with known increases and decreases in light intensity: cluster 1 is dominated by feedforward inhibitory input and cluster 3 by crossover inhibitory input.

As expected from the stimulus-dependence of feedforward and crossover inhibitory input (e.g. Figure 3A), responses in clusters 1 and 3 corresponded to different stimulus properties. Image patches eliciting responses in the first cluster were bright compared to the image mean (closed circles in Figure 3C), whereas those in the third cluster had a mean intensity similar to that of the entire image (open circles in Figure 3C). The distinctions among clusters held across cells and across images (Figure 3C, bottom). We did not probe image patches with large negative intensity because we focused on patches that elicited an increase in spike rate (as in Figure 2). What is clear from this analysis, however, is that patches with little change in mean intensity can elicit large and spatially-nonlinear crossover inhibitory input.

Excitatory responses to the image patches corresponding to each inhibitory cluster differed systematically. Excitatory responses to patches corresponding to the first cluster were large and exhibited minimal spatial nonlinearity (Figure 3D, top shows the same cell as Figure 3B). Excitatory responses corresponding to the third cluster were relatively small and exhibited strong spatial nonlinearities (Figure 3D, open circles). Across cells, the NLIs for excitatory responses to image patches corresponding to cluster 3 were larger than those for patches corresponding to cluster 1 (Figure 3D, bottom).

In summary, clustering allowed us to identify the components of inhibitory input elicited by specific natural image patches. Crossover inhibitory input showed strongly nonlinear responses to spatial structure, and was elicited by image patches that also elicited strongly nonlinear excitatory inputs. These observations suggest that crossover inhibitory input cancels nonlinear excitatory input and linearizes responses to natural inputs.

### Functional properties of inhibitory input required to cancel nonlinear excitatory input

The analyses described above indicate that crossover inhibitory input cancels nonlinear excitatory input to linearize On parasol spike responses to natural images. What functional properties of crossover inhibitory input are required for effective cancellation of nonlinear excitatory input? It is difficult to selectively manipulate inhibitory input experimentally to answer this question due to off-target effects of pharmacological agents. Instead, we used subunit receptive field models with parallel excitatory and inhibitory paths (Figure 4A; see Methods); these are an extension of the bipolar cell subunit receptive field models we used previously to account for Off parasol responses to natural images (Turner and Rieke, 2016). These models allowed us to investigate how sensitivity to spatial structure in the input depended on specific properties of the inhibitory circuitry.

**Figure 4:**
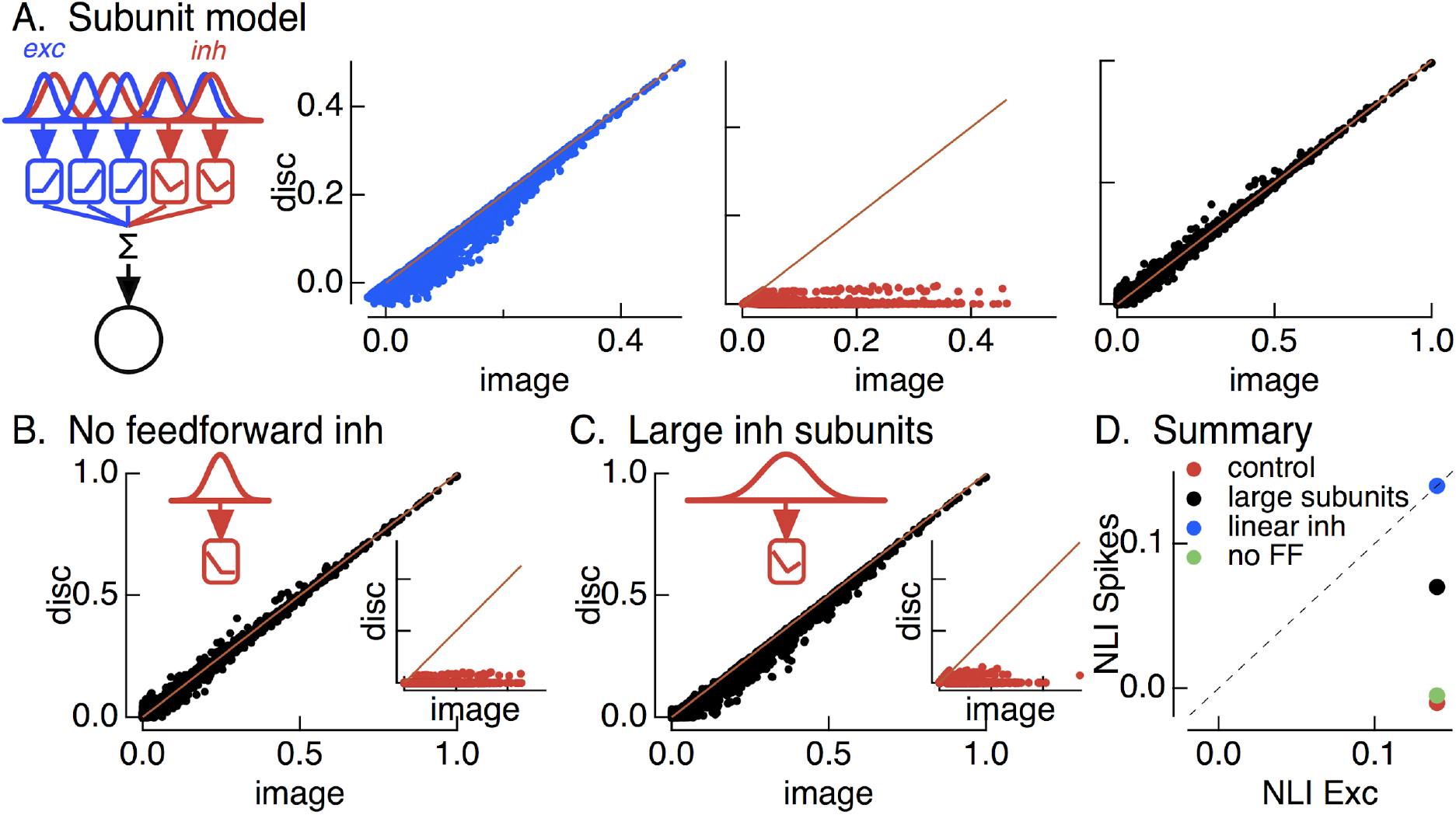
Crossover inhibitory synaptic input is necessary and sufficient for linear spatial integration. A. Model construction. (left) Architecture of subunit models. Parallel pathways generated excitatory (blue) and inhibitory (red) synaptic input to a RGC. Each pathway was composed of multiple subunits, each with a separate spatial filter and output nonlinearity. Excitatory and inhibitory subunits covered the same region of space but were positioned independently. (right) Excitatory inputs (blue), inhibitory inputs (red) and spike outputs (black) for ‘default’ model parameters (see Methods). B. Spike output for model lacking feedforward inhibitory synaptic input. Feedforward inhibitory input was eliminated by changing the shape of the nonlinearity in the inhibitory pathway (inset). C. Spike output for model in which spatial filters for inhibitory subunits are doubled in size and density of inhibitory subunits is correspondingly decreased. D. Summary of difference in NLI for spikes and excitatory input for several models for inhibitory input (see also Figure 2E).

The first stage of the model filters spatial input through two regularly-spaced grids of subunits -- with separate subunits for excitatory and inhibitory paths. The receptive field size of each subunit was set to be consistent with responses to contrast-reversing gratings (Turner and Rieke, 2016). The filtered signal in each subunit was passed through a nonlinearity determined by the contrast-response function for either excitatory or inhibitory synaptic input (Figure 4 - Figure Supplement 1). Inhibitory subunits had “U” shaped nonlinearities, reflecting both feedforward and crossover components. Excitatory (blue) and inhibitory (red) subunit outputs were weighted by a Gaussian profile representing the receptive field center and then summed, with a three-fold larger weighting of excitatory inputs reflecting the larger driving force near spike threshold. Spike responses (black) were predicted by thresholding this summed signal to eliminate negative responses. Figure 4A shows modeled excitatory, inhibitory and spike responses for a collection of natural image patches from a single image. This ‘base’ model captured the measured cancellation of nonlinear excitatory input by inhibitory input and the linearity of the spike response (Figure 4A, right); consistent with experiment (Figure 2D), the NLI for excitatory input was larger than that for spike output (Figure 4D, red). We did not attempt to make quantitative predictions of responses to specific patches because of uncertainties in model details such as the subunit positions in a recorded cell.

We next probed how three manipulations of inhibitory input affected model responses. First, we replaced the measured inhibitory subunit nonlinearity with a linear contrast-response function. This forces inhibitory input to integrate linearly across space. Inhibitory input in this case was unable to cancel nonlinear excitatory input, and the NLIs for excitatory input and spike output were similar (Figure 4D, blue). This result highlights the necessity of nonlinear spatial integration if inhibitory input is to cancel nonlinearities in excitatory input. Second, we eliminated feedforward inhibitory input by setting the positive-contrast region of the contrast-response function for inhibitory input to zero (Figure 4B). This had minimal effect on the model spike response (Figure 4D, compare green and red). Hence, crossover inhibitory input alone was sufficient to cancel nonlinear excitatory input, and feedforward inhibitory input contributed little to shaping model responses (see also (Cafaro and Rieke, 2013). Third, we doubled the size of the inhibitory subunits without changing the size of the excitatory subunits (Figure 4C). This substantially decreased the ability of inhibitory input to cancel nonlinear excitatory input (Figure 4D, compare black and red). This analysis suggests that linearity of the spike response requires that nonlinear subunits for excitatory and inhibitory circuits have similar sizes.

In summary, the modeling of Figure 4 indicates that the insensitivity of the On parasol spike response to spatial structure in natural inputs depends on nonlinear crossover inhibitory input that has a similar spatial scale as nonlinear excitatory input. Importantly for the analyses below, this means that Off retinal circuits are the main source of the inhibitory input that regulates spatial integration.

### Spatial integration of flashed vs modulated gratings differs

The experiments described thus far (1) indicate that the linearity of On parasol responses to spatial structure in natural images originates from recruitment of inhibitory input, and (2) identify key properties of inhibitory input that make it effective in linearizing spike responses. This description, however, does not explain why On parasol cells are sensitive to spatial structure in contrast-reversing gratings but not in natural images (Figure 1). The experiments and analyses described in the next two sections indicate that the periodic time course of the grating stimuli substantially impacts synaptic integration by modulating the adaptational state of the cone photoreceptors. This time-dependent adaptation contributes strongly to cycle-average responses to contrast-reversing gratings, such as those in Figure 2G and many other studies.

We start by analyzing On parasol responses to flashed gratings (Figure 5A, B), since these can be compared directly to the flashed images in Figure 2. As shown in Figure 1C, the onset of a flashed grating elicited a minimal spike response, unlike the grating offset which elicited a clear response. This difference held across cells (Figure 5B, left; onset response 0 ± 5 spikes, offset response 57 ± 7 spikes, mean ± sem, n=7, p < 0.001). To explore the origin of this difference, we compared excitatory and inhibitory inputs produced at grating onset and offset (Figure 5A). Both the relative magnitude and the timing of excitatory and inhibitory inputs differed for grating onset and offset: the I/E ratio was ~2-fold larger at grating onset than offset (Figure 5B, middle; p < 0.01), and excitatory input was delayed relative to inhibitory input at grating onset but not offset (Figure 5B, right; onset p < 0.01, offset p = 0.9). Both of these factors should contribute to the larger spike response at grating offset.

**Figure 5:**
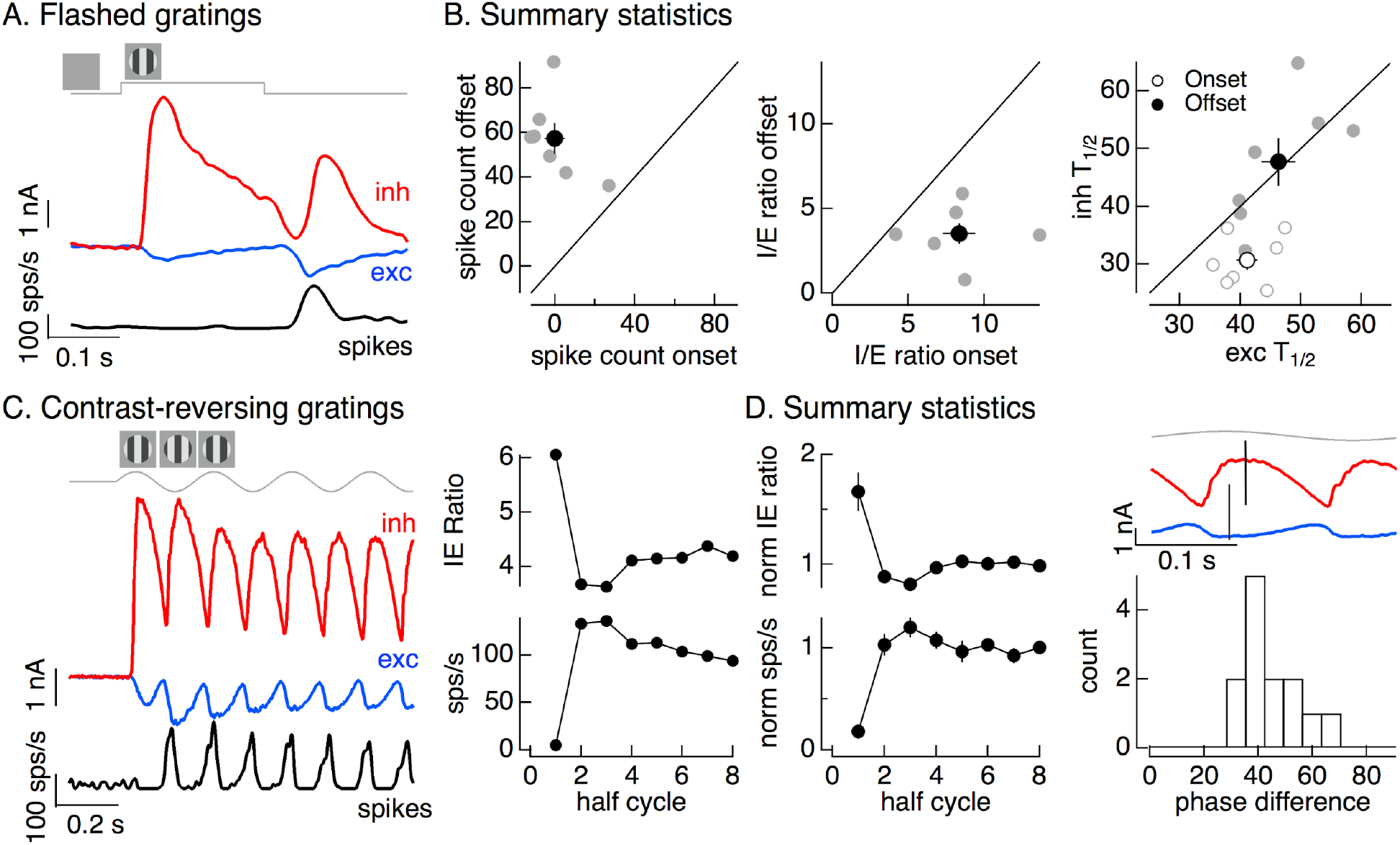
Time dependence of On parasol responses to flashed and contrast reversing gratings. A. Excitatory and inhibitory synaptic inputs and spike response to a flashed grating. B. Summary of responses to flashed gratings from 7 On parasol RGCs. (left) Spike count at grating offset vs that at grating onset. (middle) Ratio of inhibitory input to excitatory input at grating offset vs that at grating onset. (right) Relative timing of excitatory and inhibitory synaptic inputs, measured as the time to reach half-maximal amplitude. C. (left) Excitatory and inhibitory synaptic inputs and spike output in response to a contrast-reversing grating modulated at 4 Hz. (right) I/E ratio and spike count for each grating half-cycle. D. Summary of statistics of grating responses across 13 On parasol RGCs. (left) I/E ratio and spike count have been normalized by the mean value for a cycle number of 6 or more. (right) Histogram of time of peak excitatory input relative to inhibitory input. Response timing estimated by fitting each response with a sinusoid. For 4 Hz stimuli, a phase difference of 40 degrees corresponds to 28 ms.

The difference in responses at the onset and offset of a flashed grating suggested that responses to contrast-reversing gratings might also depend on time since grating onset. This was indeed the case (Figure 5C, left). Specifically, the spike response during the first half cycle of a contrast-reversing grating was substantially smaller than that during subsequent half cycles (Figure 5C, left and bottom right). Similarly, the ratio of inhibitory input to excitatory input was substantially larger during the first half cycle of the grating than during subsequent half cycles (Figure 5C, left and top right). These features held across cells (Figure 5D, left). The kinetics of excitatory and inhibitory synaptic inputs in response to a modulated grating also differed, with increases in excitatory inputs leading increases in inhibitory inputs (Figure 5D, right). The firing rate was highest in the time window in which excitatory inputs had increased but inhibitory input was still small (e.g. compare black and blue traces in Figure 5C). Off parasol cells exhibit the opposite behavior, with the smallest I/E ratios and largest spike responses during the first half cycle of a grating (Figure 5 - Figure Supplement 1).

These analyses show that On parasol cells have minimal responses to the onset of a flashed grating and to the first half cycle of a contrast-reversing grating, indicating linear or near-linear spatial integration. The cells respond strongly at the offset of a flashed grating or in later cycles of a contrast-reversing grating, indicating the emergence of nonlinear spatial integration. This time dependence can go un-noticed in cycle-averaged responses to modulated gratings (e.g. Figure 1A; Turner and Rieke, 2016).

### Cone adaptation shapes synaptic integration

The time course of the light intensity encountered by a single receptive-field subunit differs substantially between the onset and offset of a flashed grating and likewise between the first and subsequent half cycles of a modulated grating. Prior to the onset of either stimulus type, subunits encounter the mean stimulus intensity, while at later time points they undergo larger dark-light or light-dark transitions. Several nonlinear mechanisms could cause this difference in stimulus time course to contribute to the time-dependent responses illustrated in Figure 5. One such mechanism is adaptation in the cone photoreceptors, which we explore below.

We used an empirically-derived biophysical model of cone phototransduction to predict outer segment current responses to contrast-reversing gratings (see Methods and Angueyra et al., 2021 for details). Modeled cone photocurrents showed three features that could potentially shape RGC responses (Figure 6A): (1) the amplitude of the cone response was smaller on the first half cycle of the grating than on subsequent cycles; (2) cone responses were asymmetric, with larger responses to the dark bars of the grating (decrements) than to bright bars (increments); and, (3) the kinetics cone responses to dark-to-light transitions were sped relative to responses to light-to-dark transitions. These features are shared by responses of real cones (Figure 6 - Figure Supplement 1 and Angueyra et al., 2021).

**Figure 6:**
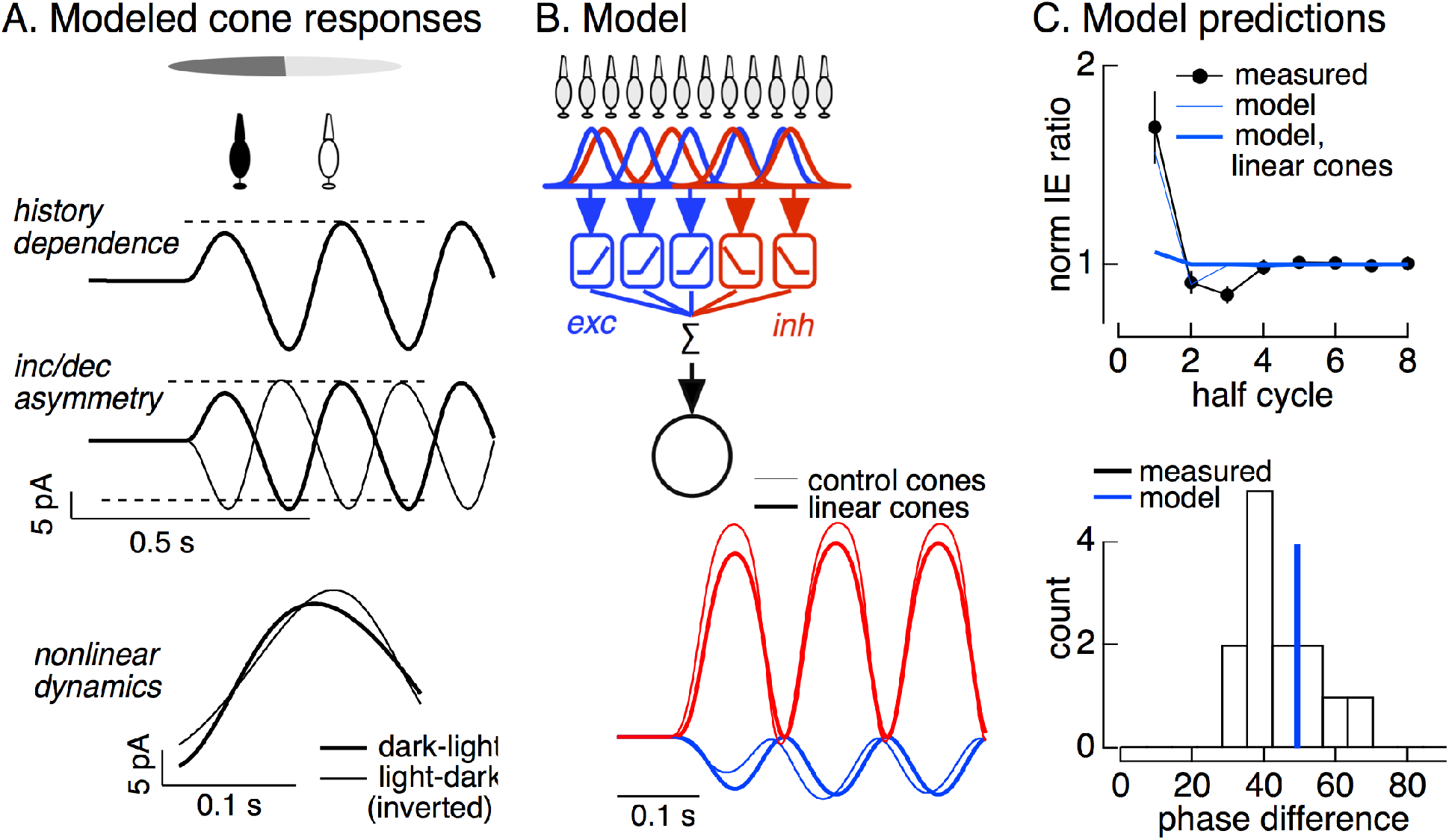
Nonlinearities in cone responses to contrast-reversing gratings and implications for On parasol synaptic inputs. A. A quantitative model for cone phototransduction currents predicts three properties of cone responses to contrast-reversing gratings that might shape time course of RGC responses: (1) a dependence on stimulus history, (2) an asymmetry between light increments and decrements, and (3) a difference in kinetics of cones responding to light-to-dark vs dark-to-light transitions. B. Circuit model to explore how properties of cone responses from A could alter RGC responses. The model from Figure 4 was adapted to incorporate a first stage based on models for the cone responses. Subunit nonlinearities were set by measured contrast-response functions. Shown are examples of excitatory and inhibitory synaptic inputs predicted by models with adapting and linear cone models. C. Comparison of model predictions with experiment. (top) Histogram of I/E ratio from On parasol recordings (from Figure 5D) and predictions from models with adapting (thin blue line) and non-adapting (thick blue line) cones. (bottom) Histogram of phase difference between excitatory and inhibitory synaptic inputs elicited by contrast-reversing gratings (from Figure 5D) and prediction from model (blue vertical line).

Different aspects of light stimuli control excitatory and inhibitory synaptic inputs to On parasol cells: light increments dominate excitatory inputs, whereas light decrements dominate inhibitory inputs (Figure 3A). This suggests that the asymmetry in the amplitude and timing of increment and decrement responses in the cones could control the relative magnitude and timing of excitatory and inhibitory synaptic inputs to an On parasol RGC and hence how these inputs are integrated to control spike output. The difference in the time course of responses of On and Off parasol cells to the onset of a grating (compare Figure 5C and Figure 5 - Figure Supplement 1) supports this suggestion.

To test for a role of cone adaptation in controlling spatial integration, we modified the subunit model from Figure 4 to take cone signals as input (see Methods). Hence, we simulated responses of an array of cones to contrast-reversing gratings, and then used these simulated cone responses as input to models with excitatory and inhibitory subunits, including measured contrast-response functions. We used this approach to predict responses to gratings for models incorporating both adapting and non-adapting (i.e. linear) cones (Figure 6B; Figure 6 - Figure Supplement 2). Note that the measured RGC contrast-response functions should capture time-independent (i.e. static) nonlinearities in the cone responses, particularly the asymmetry between the amplitude of increment and decrement responses; these contrast-response functions, however, will not capture time-dependent nonlinearities in the cone responses.

Models with adapting cones predicted a 70-80% larger I/E ratio on the first grating cycle compared to the steady-state ratio (Figure 6C, top, thin blue line). This time-dependence of the I/E ratio was very similar to that observed in RGC synaptic inputs (black) and was absent in predictions of models with linear cones (thick blue line). Models with adapting cones also captured the temporal delay between excitatory and inhibitory input (thin lines in Figure 6B; blue line in Figure 6C, bottom); this delay was again absent for models with linear cones (thick lines in Figure 6B). Excitatory input is dominated by cones undergoing a dark-to-light transition, whereas inhibitory input is dominated by cones undergoing a light-to-dark transition. The timing difference in the model originates from the temporal asymmetry in the responses of cones undergoing light-to-dark and dark-to-light transitions (Figure 6A, bottom). This difference in cone kinetics is a property of adaptation and is absent in linear models of the cone responses. Thus, adaptation shapes the dynamics of the cone responses in a manner that can explain the spatially-nonlinear responses to a temporally-modulated grating.

### Removing nonlinearities in cone responses reduces nonlinear spatial integration

To directly test the hypothesis that cone adaptation shapes On parasol RGC responses to contrast-reversing gratings, we used the cone model to design grating-like stimuli that minimized nonlinearities in the cone response; we refer to this procedure as the “light-adaptation clamp” (see Methods). Specifically, we transformed a grating stimulus to minimize the difference between the response of the adapting cone model to the transformed stimulus and the response of a linear cone model to the original stimulus (Figure 7A). In the case of sinusoidal stimuli such as gratings, this procedure means identifying a stimulus for which the predicted cone response is sinusoidal, since a linear cone model predicts a sinusoidal response to a sinusoidal input. Cone responses to these modified grating stimuli are predicted to exhibit less initial history dependence and symmetrical responses to increments and decrements (in both dynamics and amplitude). We tested these stimuli in recordings from voltage-clamped cones (Figure 7B, top). Across recorded cones, the transformed stimuli altered cone responses as predicted by the model (Figure 7B, bottom).

**Figure 7:**
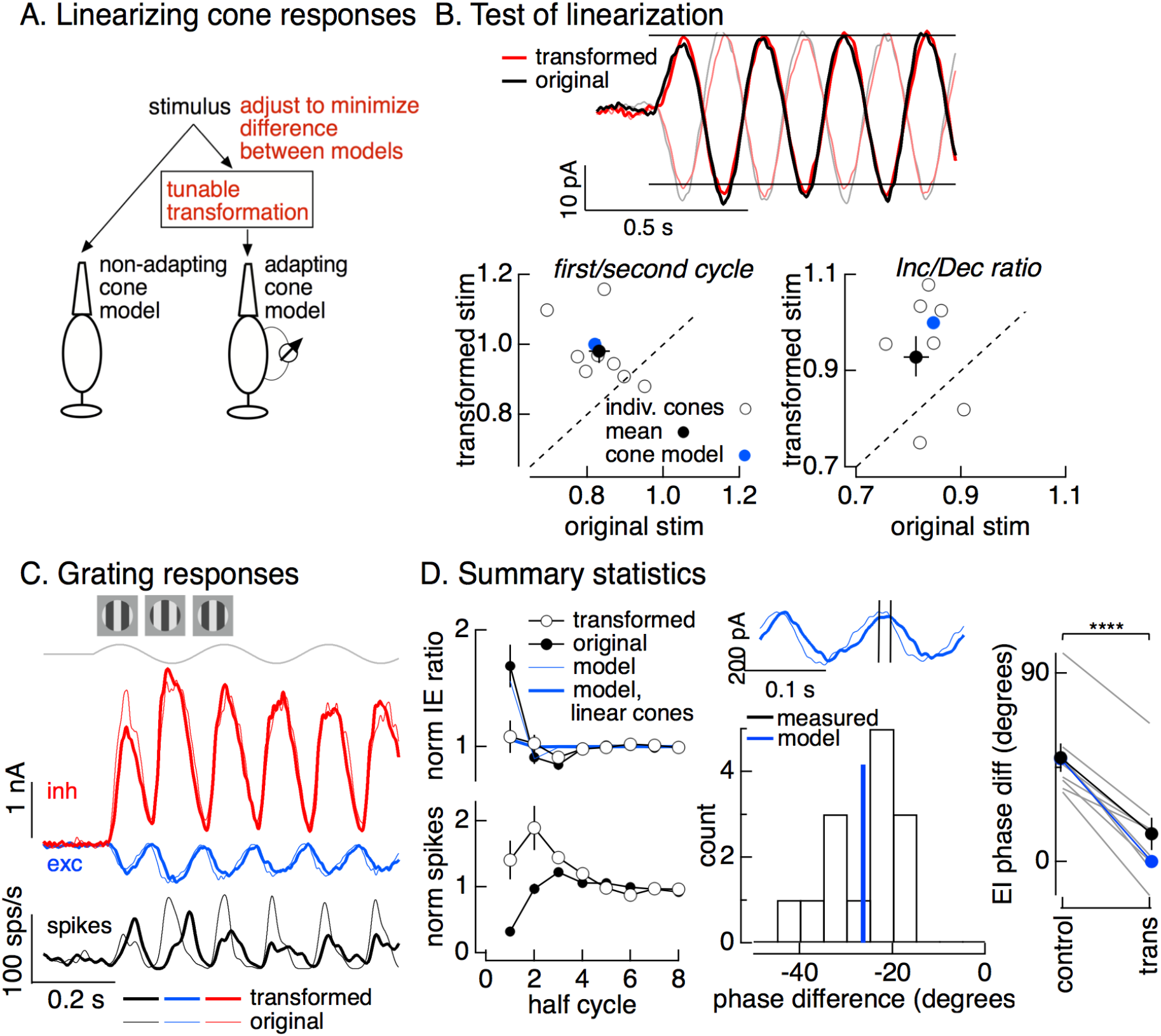
Minimizing cone adaptation minimizes time dependence of On parasol responses to contrast-reversing gratings. A. Approach to minimize nonlinearities in cone responses. Standard grating stimuli were transformed to minimize the difference between the response of an adapting cone responding to the transformed grating and a linear (non-adapting) cone responding to the original grating. B. Test of procedure from A. (top) Measured cone responses to original (black and gray) and transformed (red) grating stimuli. (bottom) Summary across recorded cones. Responses to the transformed stimulus showed less history dependence (left; measured from the ratio of the amplitude of the response on the first cycle of the grating to that on the second cycle) and less of an increment/decrement asymmetry (right). C. On parasol responses to original (thin traces) and transformed (thick traces) grating stimulus identical to those used in Figure 6. D. Summary of responses to original and transformed gratings from 9 On parasol cells. (left) I/E ratios and spike counts have been normalized in each cell by the responses for grating cycles > 6. (middle) Histogram of difference in timing of excitatory input in response to original and transformed gratings. A phase shift of 30 degrees corresponds to a 21 ms difference in timing. (right) Difference in timing of excitatory and inhibitory synaptic inputs for original and transformed gratings and prediction from model of Figure 6B (blue).

If adaptation in the cones contributes to the small spike response of On parasol RGCs during the first half cycle of a contrast-reversing grating (Figure 5C,D), then On parasol responses to the modified stimuli should show less time dependence. This was indeed the case (Figure 7C, thick traces are responses to the transformed stimuli and thin traces are responses to the original grating). Across cells, minimizing the nonlinearities in the cone responses decreased the time dependence of the I/E ratio and the spike response (Figure 7D, left). The I/E ratios for both original and transformed stimuli agreed well with those predicted from the models of Figure 6B with nonlinear or linear cone inputs (blue traces in Figure 7D, top left). Excitatory inputs in response to the transformed stimuli were substantially delayed compared to excitatory inputs in response to the original stimuli (Figure 7D, middle). This shift in timing of excitatory input caused it to overlap more with inhibitory input, and the substantial phase shift between excitatory and inhibitory input observed for conventional gratings was almost entirely eliminated by the transformed gratings (Figure 7D, right). The difference in timing of excitatory and inhibitory inputs for both grating types was well predicted by the simple model of Figure 6B (blue line and points in Figure 7D, right); this suggests that the temporal differences in cones exposed to increments and decrements (Figure 6A) plays a central role in determining the kinetics of excitatory and inhibitory inputs.

The combination of modeling in Figure 6 and direct manipulation of the cone responses in Figure 7 indicates that cone adaptation can explain the majority of the observed time-dependence of the On parasol responses to contrast-reversing gratings. The decrease in I/E ratio after grating onset and the speeding of excitatory input relative to inhibitory input account for the strength of the steady-state responses to contrast-reversing gratings, and both of these depend on adaptation in the cones.

### Cone adaptation and circuit nonlinearities can explain linear responses to natural images

Natural inputs vary over time due to eye movements and object motion within an image. Hence it is not immediately clear whether the linear spatial integration that On parasol cells exhibit at grating onset or the nonlinear integration during later grating cycles is more relevant for natural images. To distinguish between these possibilities, we used the cone/subunit model described above to predict how cone adaptation impacts the I/E ratio and sensitivity to spatial structure of On parasol cell responses to time-varying natural inputs.

Stimuli were created by sampling a static natural image with a simulated eye movement trajectory. These stimuli are identical to the movies used in Figure 1 except for the use of simulated rather than real eye movements. Eye movements were simulated by a diffusional process to account for fixational eye movements, interrupted by occasional large and discrete changes in position to account for saccades (see Methods for details). The resulting spatio-temporal pattern of inputs was converted to signals in the modeled cone photoreceptor array (Figure 8A, bottom, shows modeled responses of a single cone). Adaptation substantially attenuated the amplitude of modeled cone responses compared to those without adaptation (Figure 8A, bottom).

**Figure 8:**
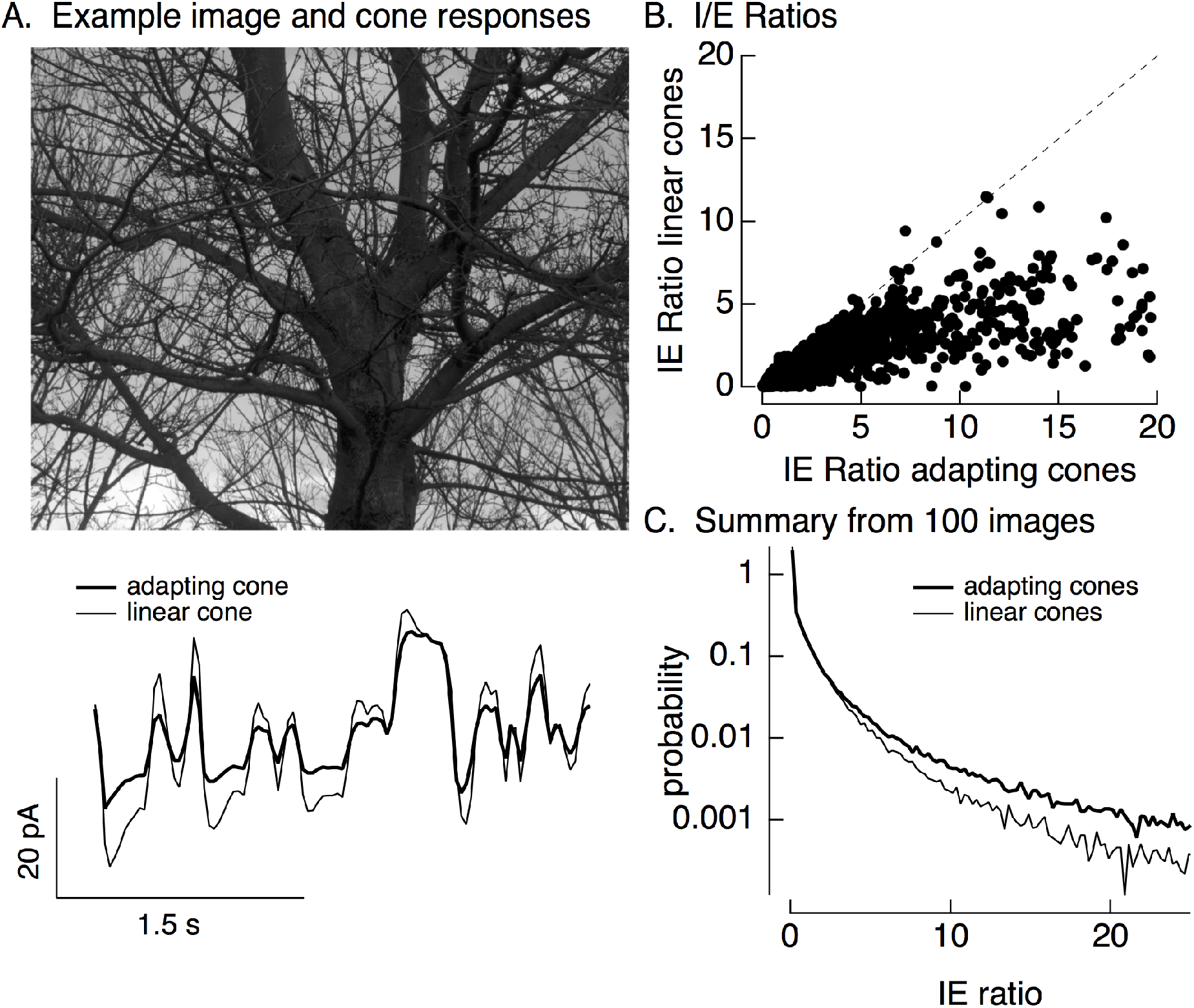
Cone adaptation substantially increases I/E ratio for natural images. The model from Figure 6B with either adapting or non-adapting cones was used to predict excitatory and inhibitory synaptic inputs during eye movements about a scene (A). Cone responses are shown below the image. I/E ratios were computed for every 30 ms time window (B). Ratios for linear cones were systematically smaller than those for adapting cones. This was consistent across images, as indicated by histograms of I/E ratios across many images (C).

We used the modeled adapting and non-adapting cone signals as input to subunit models for excitatory and inhibitory synaptic inputs as in Figure 6B. We used these models to predict I/E ratios in every 30 ms time bin (Figure 8B). I/E ratios varied widely across time bins, but were systematically larger for models with adapting cones compared to models with non-adapting cones (Figure 8B). To test for a similar effect across images and trajectories, we constructed histograms of I/E ratios for models with adapting and non-adapting cones; Figure 8C compares histograms from many images, each with an independent eye movement trajectory. Cone adaptation substantially increased the number of patches in which inhibitory input was sufficiently large to attenuate excitatory input (I/E ratios exceeding 3).

This analysis suggests that cone adaptation shapes responses to contrast-reversing gratings and time-varying natural inputs quite differently. This difference can be explained by the different luminance trajectories encountered by cones during these stimuli, and the corresponding differences in the time course of cone adaptation. In the case of a sinusoidally-modulated grating, an increment in intensity at a given spatial location always occurs after a decrement, and likewise decrements occur after increments. Hence a cone encountering an increment starts from a state in which signaling gain is high (due to the preceding decrement and relief of adaptation), and a cone encountering a decrement starts from a low gain state (due to the preceding increment and engagement of adaptation). This both speeds and increases the amplitude of responses of cones signaling increments relative to those signaling decrements - which in the case of an On parasol cell means shifting signaling in favor of excitatory synaptic inputs over inhibitory synaptic inputs (see Figure 6). Naturalistic inputs typically do not show such predictable behavior: increments are not systematically preceded by low intensity regions of a scene. Instead, on average, cones encountering a light increment or decrement start from the average image intensity - much like the first half-cycle of a modulated grating or the onset of a flashed grating. In all such cases, cone adaptation had a strong impact on the I/E ratio and spike response (e.g. linear vs adapting cones in Figures 6 and 8 and control vs transformed stimuli in Figure 7). Specifically, adaptation in the cones shifted the I/E balance in favor of inhibitory input, and this in turn decreased the sensitivity of On parasol RGC responses to spatial structure.

### Unique natural stimulus features coded by coactivation of ON and OFF cells

On and Off ganglion cells of the same type are classically assumed to respond to opposite polarity but otherwise identical stimulus features. Asymmetries between On and Off cells challenge this simple picture (Chichilnisky and Kalmar, 2002; Zaghloul et al., 2003; Sagdullaev and McCall, 2005; Murphy and Rieke, 2006; Nichols et al., 2013; Turner and Rieke, 2016; Ravi et al., 2018). The experiments and analysis below suggest that asymmetries in spatial integration (as in Figure 1) cause joint activation of On and Off parasol cells to encode image features not encoded by either cell type alone. This analysis and the associated conclusions are restricted to activation of the receptive field center.

We used subunit models to identify image patches predicted to coactivate On and Off parasol cells. We first sampled patches of natural inputs through a grid of subunits as in Figure 4. We then constructed two RGC models (see Methods and (Turner and Rieke, 2016): (1) in a spatially-linear model, subunit outputs were weighted by a Gaussian receptive field with a size approximating the parasol receptive field center and then linearly summed; (2) in a spatially-nonlinear model, subunit outputs were rectified, weighted by a Gaussian and summed. Unlike the model of Figure 4, these models do not incorporate separate excitatory and inhibitory paths, but they do capture the linear vs nonlinear difference in how On and Off parasol RGCs integrate inputs across space.

We used responses to many image patches to define constant-response contours for each of these models in a space spanned by the local luminance and spatial contrast of the sampled patches (patch statistics were measured within the receptive field center). These contours identify the set of image properties (i.e. local luminance and spatial contrast) that resulted in similar responses of a given RGC model and hence that cannot be distinguished based on the model response.

As expected, response contours for spatially-linear models (On parasol and linear Off parasol) do not depend on spatial contrast (i.e. the contours are near horizontal; Figure 9A, left and middle). The contours for the spatially-nonlinear Off parasol cell, however, do depend on spatial contrast because of nonlinear spatial integration (Figure 9A, right). This dependence means that image patches with positive local luminance and high spatial contrast should activate both On and Off parasol cells; such coactivation would be absent if both cells exhibited linear spatial integration. These patches are located at the bright side of edges in an image (Figure 9B).

**Figure 9:**
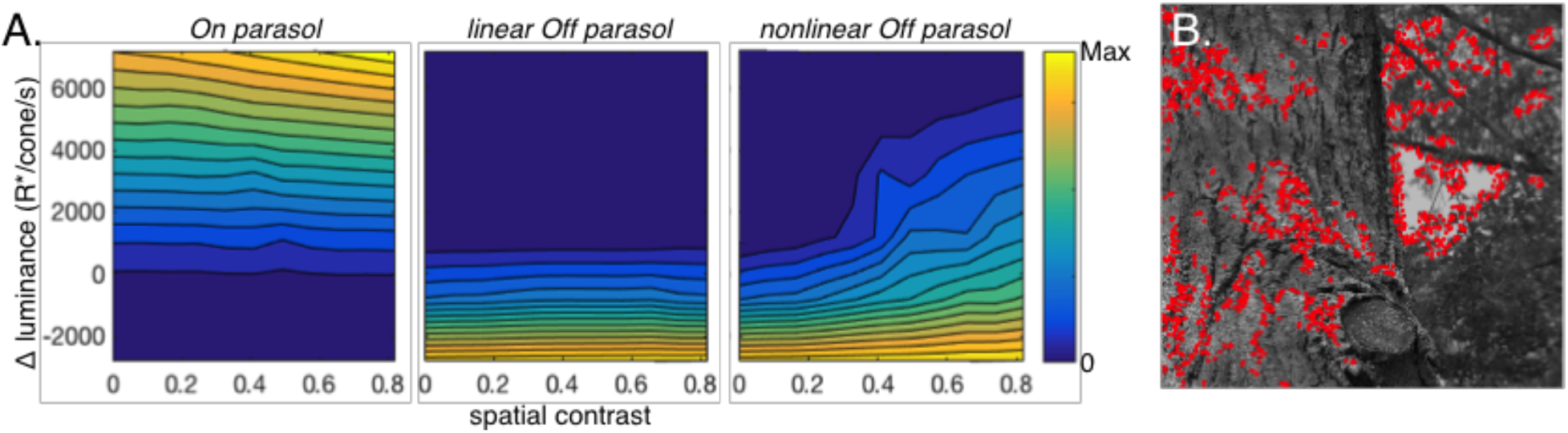
Joint activity of On and Off parasol RGCs encodes patches with positive luminance and high spatial structure. A. Responses of On and Off parasol models as a function of the spatial contrast (x-axis) and change in mean luminance (y-axis) for a collection of image patches. Panels show constant-response contours for On parasol cells (left), Off parasol cells with linear spatial integration (middle), and Off parasol cells with nonlinear spatial integration (right). B. Image patches eliciting activity of both On and Off parasol cells for nonlinear Off parasol model. These are in the upper right corner of the contour plots in A.

The modeled response contours of On and nonlinear Off parasol cells have different dependencies on changes in mean luminance and contrast for image patches in which the cells are coactive: On parasol model responses are affected only by changes in luminance, while Off parasol model responses depend on both luminance and spatial contrast. This means that the joint responses of On and Off cells should permit discrimination of image patches with small differences in local luminance and/or spatial contrast (Figure 9 - Figure Supplement 1); such discrimination is not possible from responses of either cell type considered alone. Consistent with this modeling, image patches with similar mean luminance within the receptive field center that elicited near-identical measured spike responses from On parasol cells could elicit quite different responses from Off parasol cells (Figure 9 - Figure Supplement 2).

This analysis indicates that joint activity of On and Off parasol cells may encode stimulus features that would be ambiguous if both cell types integrated linearly over space. Note that our analysis here focuses entirely on stimuli restricted to the receptive field center. The improved coding of specific stimulus aspects comes with a cost of sensitivity to other stimulus features. For example, with nonlinear spatial integration, Off parasol cells do not uniquely encode decreases in luminance. We return to this issue in the Discussion.

## Discussion

The manner in which retinal ganglion cells integrate inputs across space has important functional and mechanistic implications. Mechanistically, the linearity of spatial integration is used to classify RGC types and gives insight into the operation of the upstream circuits that provide input to the RGC (Enroth-Cugell and Robson, 1966; Hochstein and Shapley, 1976,; Demb et al., 1999). Functionally, nonlinear spatial integration enhances RGC sensitivity to some features of the input at the expense of others (Schwartz et al., 2012) and is a central component in many specialized retinal computations (reviewed in Gollisch and Meister, 2010). Here, we show that the qualitative nature of spatial integration - i.e. linear vs. nonlinear - is dictated by several nonlinear circuit mechanisms acting in concert. The interaction of these mechanisms to control spatial integration results in a striking difference in how On parasol RGCs respond to spatial structure in temporally-modulated grating stimuli compared to natural inputs.

### Neural computation and responses to complex stimuli

Artificial stimuli, by design, often activate specific circuit mechanisms while avoiding activation of other mechanisms. But complex stimuli, including natural inputs, recruit multiple circuit mechanisms that act in concert to control circuit outputs. Interactions among these coactive mechanisms pose a fundamental challenge to understanding the operation of neural circuits.

Here, we show that the manner in which On parasol RGCs respond to spatial structure in their inputs is shaped by a combination of adaptive nonlinearities in cone phototransduction together with rectifying synaptic nonlinearities in the circuits generating excitatory and inhibitory inputs. Thus, for contrast-reversing gratings, cones encountering increases and decreases in light intensity start from different adaptation states and consequently generate responses with different amplitudes and kinetics. These differences in cone inputs to On and Off circuits, coupled with rectification in the circuits generating excitatory and inhibitory input to On parasol cells, favor excitatory input over inhibitory input and contribute to strong responses to the spatial structure of the grating. Natural inputs, however, lack the periodicity of contrast-reversing gratings; as a consequence, inhibitory input is both large and temporally in phase with excitatory input, and hence is well-poised to decrease sensitivity to fine spatial structure. The opposite is true for Off parasol cells - where the periodic changes in cone adaptation during grating stimuli favor inhibitory input over excitatory input. This provides a clear example, with clear functional consequences (see below), of how several nonlinear mechanisms acting in concert control circuit outputs in unexpected ways.

The differences in cone signaling that control the linearity or nonlinearity of On parasol spatial integration are subtle. Yet these relatively small changes in input qualitatively change the computational properties of the circuit. This is one of several examples in which control of computation relies on fine tuning of common circuit elements rather than recruitment of distinct circuits or other more exotic mechanisms (see also Grimes et al., 2014). The sensitivity of neural circuits to subtle changes in input or in operating point will be important in constructing models for neural computation, including artificial neural networks that attempt to match the efficiency of real neural circuits in specific tasks.

### Balanced excitation and inhibition

Cortical neurons typically receive high rates of excitatory and inhibitory inputs which largely cancel -- a mode referred to as balanced excitation and inhibition (reviewed by Isaacson and Scanziani, 2011). The discovery of E/I balance has been an important breakthrough in our understanding how cortical circuits work, and disruption of E/I balance can underlie the aberrant neural signaling characteristic of several brain disorders (reviewed by Sutula and Dudek, 2007; Nelson and Valakh, 2015).

Signaling in circuits exhibiting E/I balance often relies on differences in timing of excitatory vs inhibitory inputs (reviewed by Isaacson and Scanziani, 2011). For example, in auditory (Wehr and Zador, 2003; Wu et al., 2008), somatosensory (Pouille and Scanziani, 2001; Swadlow, 2002; Wilent and Contreras, 2005) and visual cortices (Liu et al., 2010), the onset of a sensory stimulus creates a brief time window in which excitatory input exceeds inhibitory input; action potentials are preferentially generated in this window. Several circuit features can contribute to these timing differences and their impact on spike output, including delays in inhibitory signals relative to excitatory signals due to routing through extra synapses (Pouille and Scanziani, 2001; Gabernet et al., 2005) and the location of excitatory inputs on the dendrites and inhibitory inputs on or close to the soma of a target neuron (Stokes and Isaacson, 2010).

Here, we show that the relative timing of excitatory and inhibitory input is highly stimulus dependent. As described above, adaptation in the cones during periodic stimuli like contrast-reversing gratings creates kinetic differences in cones encountering increases and decreases in light intensity. Cones encountering these different polarity stimuli dominate input to the circuits generating excitatory and inhibitory input to On parasol RGCs, and the difference in kinetics provides a time window in which excitatory input substantially exceeds inhibitory input. For non-periodic stimuli, however, the adaptational state of the cones does not correlate strongly with the change in intensity encountered and cones responding to increases and decreases in light level respond with similar kinetics. Hence spatial structure in such stimuli creates near-simultaneous changes in excitatory and inhibitory input and little or no spike response.

The circuit features giving rise to this stimulus-dependent control of E/I balance are common but also distinctive. The distinct circuit features - notably the sensitivity of the circuits controlling excitatory and inhibitory inputs to different aspects of the circuit input (here increases and decreases in light level) - may help identify other circuits in which a similar control could be at work.

### Asymmetries between On and Off RGCs and neural coding

On and Off RGCs of the same type are classically assumed to respond to similar stimulus features with opposite polarity. Multiple findings show that this picture is too simple, and in fact that corresponding On and Off RGCs can exhibit asymmetries in addition to the polarity of their responses. This includes differences in receptive field size (Corson et al., 2000), contrast sensitivity (Zaghloul et al., 2003), rectification (Corson et al., 2000; Turner and Rieke, 2016), and balance of excitatory and inhibitory inputs (Murphy and Rieke, 2006). Some of these asymmetries - e.g. the smaller receptive fields of Off RGCs - have been suggested to be important in population coding of natural inputs (Balasubramanian and Sterling, 2009; Ratliff et al., 2010; Pandarinath et al., 2010; Karklin and Simoncelli, 2011). On/Off asymmetries also differ for different RGC types (Ravi et al., 2018), a factor that will need to be considered in efficient coding arguments.

On and Off parasol RGCs both exhibit nonlinear spatial integration in response to contrast-reversing gratings (Petrusca et al., 2007; Crook et al., 2008). Off parasol RGCs similarly respond nonlinearly to spatial structure in natural images, but unexpectedly On parasol RGCs do not. The difference in sensitivity to spatial structure in natural images suggests that certain features - specifically image patches that have similar positive luminance and differences in spatial contrast - may be distinguishable based on the joint activity of pairs of On and Off cells even when the responses of the individual cells are ambiguous. Determining whether this increased sensitivity to specific image features, at the expense of sensitivity to others, is advantageous will require considering how the entire RGC population encodes natural inputs. Nonetheless, the symmetry of On and Off parasol RGC responses to contrast-reversing gratings and asymmetry of their responses to natural inputs emphasizes the complexity of the relationship between the statistics of input stimuli and circuit nonlinearities that control important functional aspects of neural responses.

## Methods

### Tissue preparation and recording

Experimental procedures followed those described previously (Turner and Rieke, 2016; Turner et al., 2018). In brief, primate retinas were obtained through the Tissue Distribution Program of the Regional Primate Research Center at the University of Washington. Pieces of retina attached to the pigment epithelium were stored in ∼32–34°C oxygenated (95% O_2_/5% CO_2_) Ames medium (Sigma, St Louis, MO) and dark-adapted for > 1 hr. Pieces of retina were then isolated from the pigment epithelium under infrared illumination and flattened onto polyL-lysine slides. Once under the microscope, tissue was perfused with oxygenated Ames medium at a rate of ∼8 ml/min.

### Electrophysiology

Parasol RGCs were targeted for electrical recordings based on their characteristic soma size and response to a light step. Spike responses were measured in the cell-attached or extracellular configurations using electrodes filled with Ames medium. Synaptic inputs were measured in the whole-cell voltage-clamp configuration using electrodes (2-3 MΩ) filled with a solution containing (in mM): 105 Cs methanesulfonate, 10 TEA-Cl, 20 HEPES, 10 EGTA, 2 QX-314, 5 Mg-ATP, 0.5 Tris-GTP (∼280 mOsm; pH ∼7.3 with CsOH). To isolate excitatory or inhibitory synaptic input, cells were held at the estimated reversal potential for inhibitory or excitatory input of ∼−60 mV and ∼+10 mV. These voltages were adjusted for each cell to maximize isolation (see Cafaro and Rieke, 2013). Voltages have been corrected for a ~−8.5 mV liquid junction potential.

Retina sensitivity was checked before collecting data based on the contrast sensitivity of On parasol cells. All data collected came from retinas in which a 5% contrast spot modulated at 4 Hz produced at least a 20 spikes/s modulation in the On parasol spike rate. 60-70% of the preparations met this criterion.

### Stimuli

Stimuli were presented and data acquired using custom written stimulation and acquisition software packages Stage (stage-vss.github.io) and Symphony (symphony-das.github.io). Labwide acquisition packages can be found at https://github.com/Rieke-Lab/riekelab-package and protocols used in this study can be found at https://github.com/Rieke-Lab/turner-package and https://github.com/Rieke-Lab/baudin-package. Details of stimulus presentation followed previous work (Turner and Rieke, 2016; Turner et al., 2018). Unless otherwise noted, mean light levels produced 4,000 isomerizations (R*)/M or L-cone/s, 1,000 R*/S-cone/s and 9,000 R*/rod/s. Stimuli were restricted to a circular aperture with a diameter determined by a Gaussian approximation of the receptive field center (see Turner and Rieke, 2016).

Images were moved across the retina to simulate eye movements. For Figure 1, eye movements were taken directly from those measured in human observers viewing the corresponding image (Van Der Linde et al., 2009), resampled to our 60 Hz monitor refresh rate (see Turner and Rieke, 2016). For the modeling in Figure 9, we simulated eye movements using a random-walk process, with independent steps in x and y. Steps in the random walk occurred every ms, were discrete, had a size equal to the cone spacing in the model. This diffusive process was interrupted every 300 ms by a saccade-like jump in position. The discrete jumps in position consisted of random displacements in x and y drawn from a uniform distribution between −100 and +100 cone spacings.

### Clustering of inhibitory responses

Inhibitory synaptic inputs elicited by natural image patches were clustered in Figure 3. Input to the clustering algorithm consisted only of the time course of responses to all image patches sampled in recording from a given cell. The first three principal components of these combined responses provided a space for the clustering. Each individual response was projected along these three principal components, and the results were clustered using Matlab’s kmeans algorithm.

### Subunit models

Subunit models were used to explore spatial integration. The goal of these models was to explore the impact of specific circuit features on predicted sensitivity to spatial structure rather than explain the responses of a specific cell to specific images.

These models consisted of regular grids of subunits for excitatory and inhibitory pathways. Spatial inputs were filtered through the receptive field of each subunit, the filtered signals were (optionally) passed through a rectifying nonlinearity, subunit outputs were summed to generate the RGC input, and this summed signal was thresholded at 0 to generate a predicted spike output. Subunit nonlinearities were set equal to the average excitatory and inhibitory contrast-response functions (see Figure 4-Figure Supplement 1), and outputs of excitatory subunits were given a 3-fold larger weight than outputs of inhibitory subunits to reflect the larger driving force associated with excitatory inputs near spike threshold. Subunit receptive field sizes and spacing were set to 1/5th the RGC receptive field center size for excitatory subunits, and inhibitory subunits were 50% larger. These reflect the subunit sizes indicated by responses to contrast-reversing gratings (Turner and Rieke, 2016).

Several properties of the inhibitory subunits were altered from this “base” model for the analyses in Figure 4. Inhibitory subunit size and spacing was increased, the positive contrast part of the contrast-response function for inhibitory subunits was set to 0, or the subunit nonlinearity for inhibitory subunits was removed altogether, rendering the subunits linear.

A front-end model of cone phototransduction was added to the model for the analyses in Figures 6-8. This model consists of a set of differential equations that capture measured cone responses to a broad range of stimuli (Angueyra et al., 2021). All parameters of the cone model were set by previous measurements. Spatio-temporal inputs were passed through this front-end model, and the predicted cone responses provided input to subunit models constructed as above.

For the analysis of Figure 9, parasol models were simplified by removing the inhibitory pathway and making the excitatory pathway linear (for On parasol RGCs) or nonlinear (for Off parasol RGCs). These models captured the essential distinction between linear and nonlinear spatial integration.

### Cone light-adaptation clamp

Figure 7 uses stimuli designed to minimize nonlinearities in the cone responses. We used two models of the cone responses to identify these stimuli: (1) a full model of the cone responses that captures responses to a broad range of stimuli, including those invoking adaptation; (2) a linear model. The linear model was determined by the response of the full model to a brief, low contrast flash (i.e. a flash within the linear range of the full model behavior). The stimulus for the full model was a transformed version of the original stimulus, while the original stimulus (untransformed) provided input to the linear model. We then sought a stimulus transformation that minimized the difference between the outputs of the two models. For sinusoidal stimuli, such as the contrast-reversing gratings we used in Figure 7, this is particularly simple: the response of the linear model to these stimuli is also sinusoidal, and hence our procedure identifies a stimulus to the full model that creates a sinusoidal output. We refer to this as a “light-adaptation clamp” because the procedure aims to generate cone responses that lack adaptation.

We identified the appropriate stimulus transformation using a gradient-descent approach. We discretized the stimulus into time bins and then perturbed the stimulus at these discrete times. We retained perturbations that decreased the mean-squared difference between the two models’ responses (see Figure 7A) using Matlab’s fminsearch algorithm. We iterated this process while decreasing the size of the time bins until achieving a stable minimum of the mean-square difference.

### Statistics

Data, where appropriate, are plotted as mean ± sem. Reported p values are from Wilcoxon rank sum tests.

## Acknowledgments

We thank Mark Cafaro and Shellee Cunnington for excellent technical support. Tissue was provided by the Tissue Distribution Program at the Washington National Primate Research Center (WaNPRC), and we are grateful for assistance from the WaNPRC staff, especially Chris English. Qiang Chen, Greg Field, Will Grimes and Greg Schwartz provided invaluable feedback on an early draft of the manuscript. This work was supported by NIH grants F31-EY026288 (MHT) and EY028542 (FR).

---

**Figure 3 - Figure Supplement 1:**
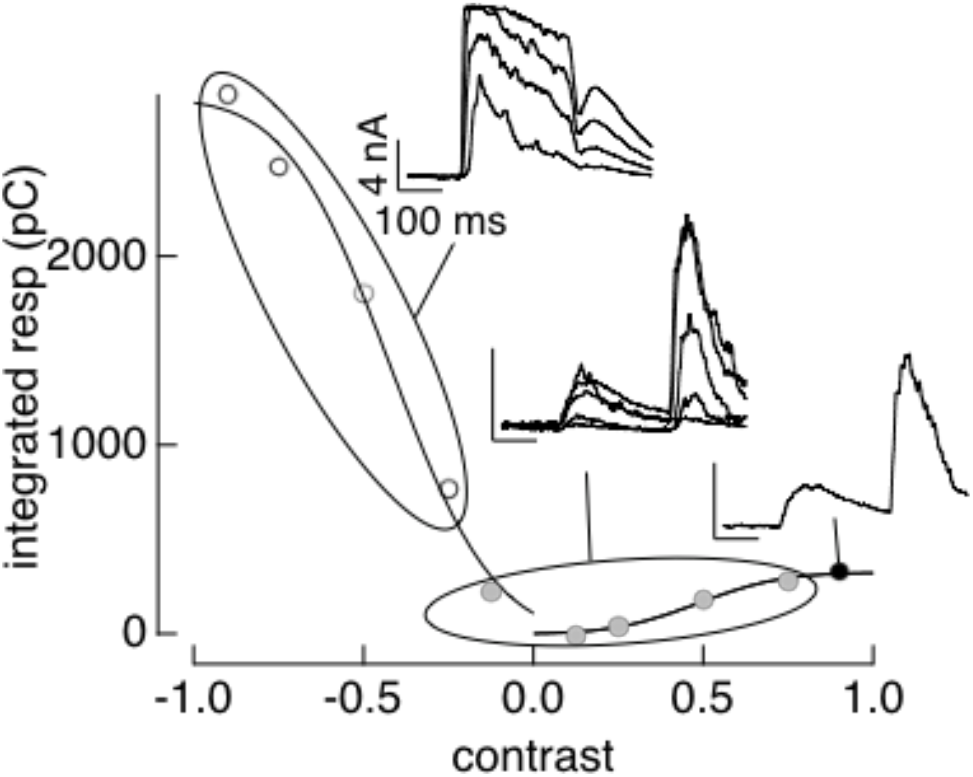
PCA-based clustering applied to contrast-response data. The cluster definitions from responses to natural image patches were applied to responses to contrast increments and decrements. Same cell as Figure 3.

**Figure 4 - Figure Supplement 1:**
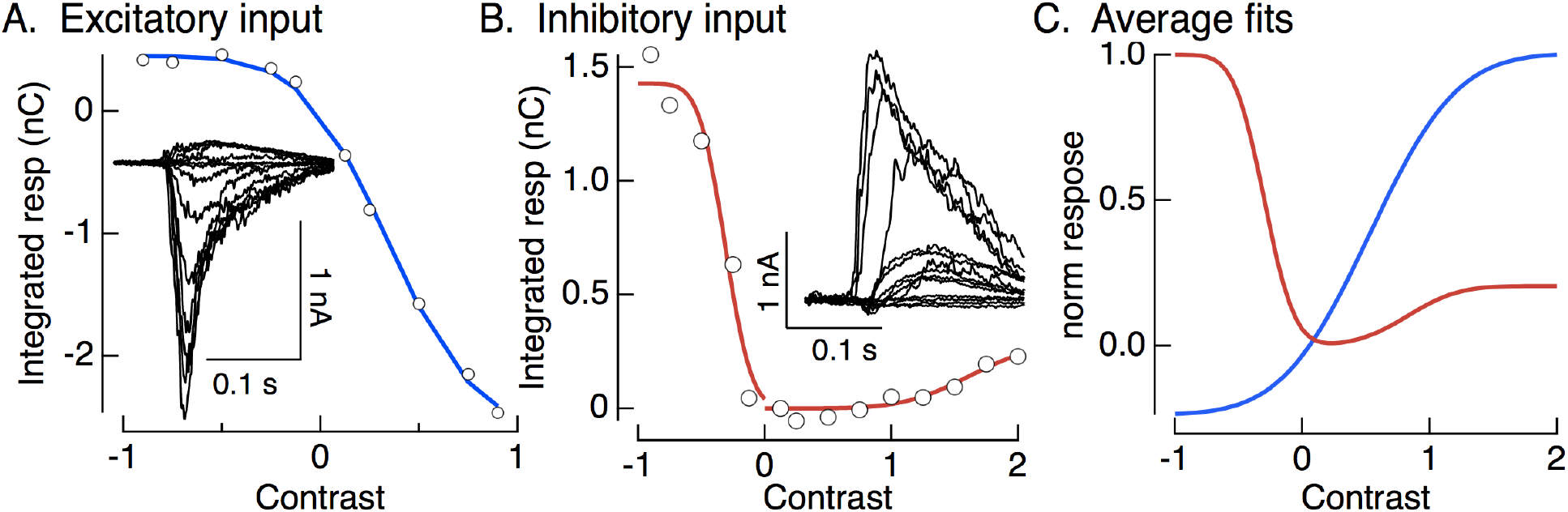
Contrast-response functions and fits. A,B. Examples of contrast-response functions for excitatory and inhibitory synaptic input to an On parasol cell. Colored smooth curves are fits (cumulative Gaussians). C. Average fits across cells. These are the nonlinearities used in the model of Figure 4A.

**Figure 5 - Figure Supplement 1:**
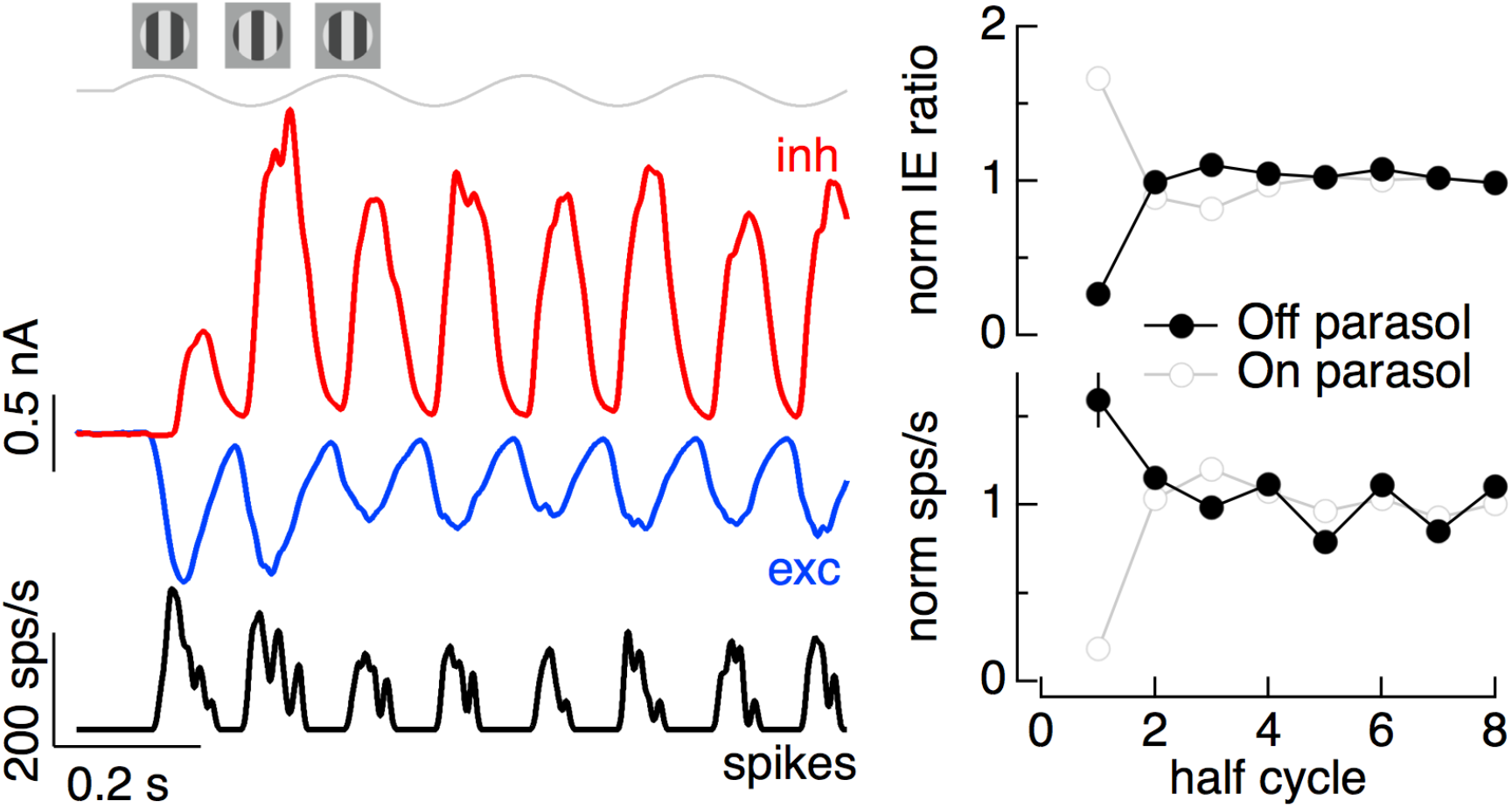
Time dependence of Off parasol responses to contrast-reversing gratings. The left panel shows the spike response and excitatory and inhibitory synaptic inputs from an example Off parasol cell for conditions identical to the On parasol recordings in Figure 5. The right panel compares the I/E ratio (top) and spike count (bottom) for 13 On and 6 Off parasol cells.

**Figure 6 - Figure Supplement 1:**
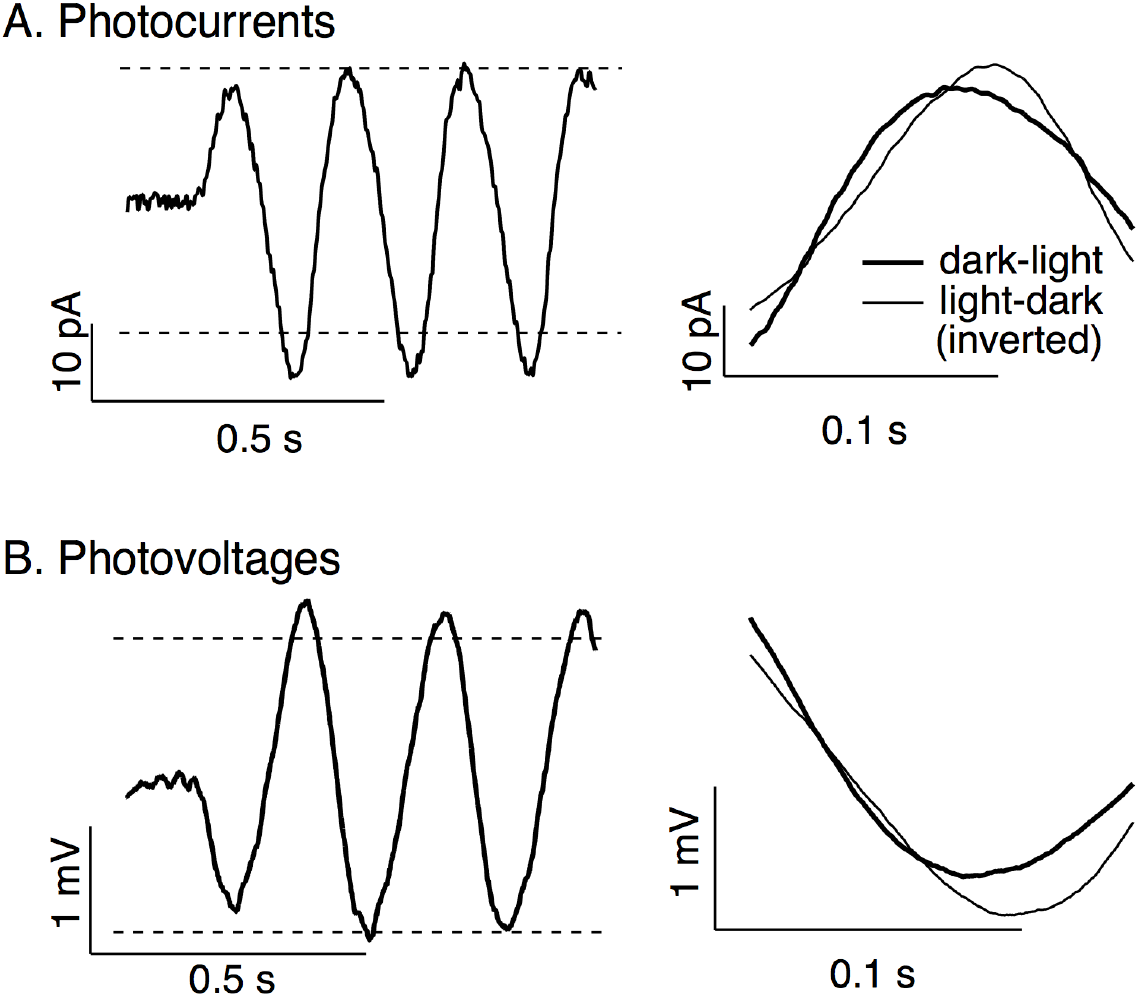
Cone photoreceptor responses to sinusoidal stimuli. Photocurrents are average responses to 5 Hz, 75% contrast sinusoidal stimuli from 6 cones. Photovoltages are average responses to 4 Hz, 50% contrast sinusoidal stimuli from 16 cones. Compare to model responses in Figure 6A. Dashed horizontal lines in left panels are displaced equally above and below the mean response to highlight the asymmetry between increment and decrement responses.

**Figure 6 - Figure Supplement 2:**
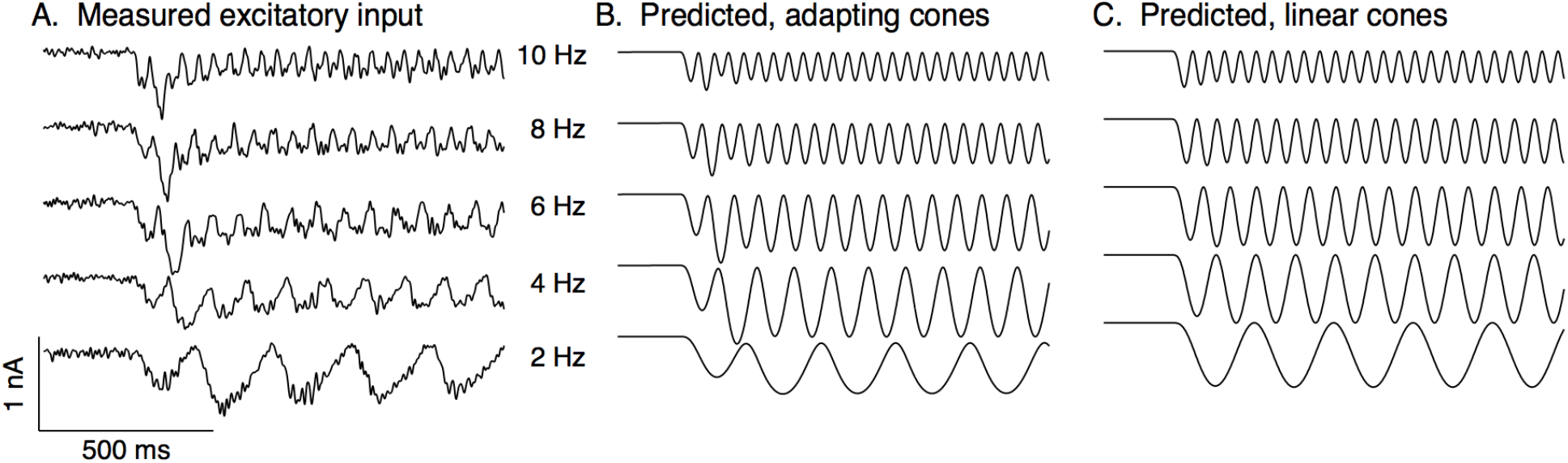
Measured and predicted excitatory synaptic inputs in responses to contrast-reversing gratings across a range of frequencies. Predictions are from the cone/subunit model in Figure 6B.

**Figure 9 - Figure Supplement 1.**
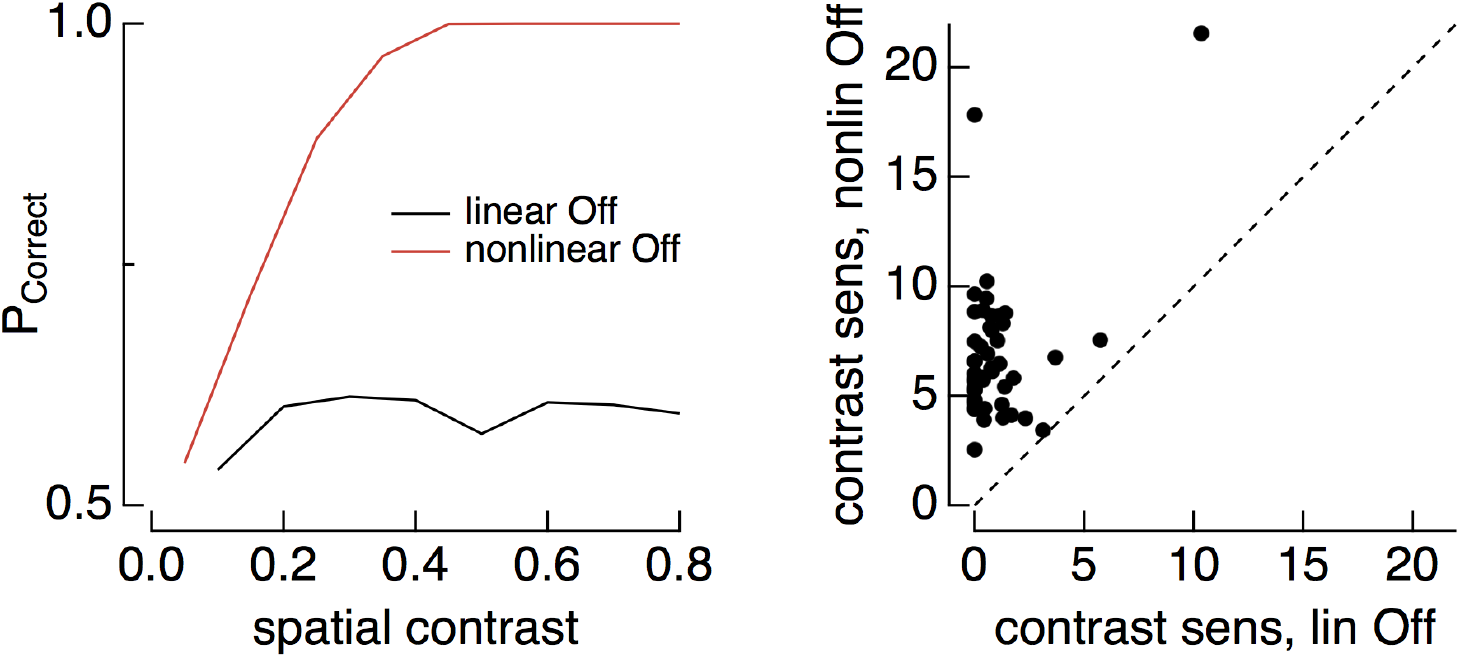
(left) Discrimination of image patches with similar mean luminance (1000 R*/cone/s) but different spatial contrast for joint responses of On and Off parasol cells. Black shows discrimination when both On and Off cells integrate linearly over space, red shows discrimination with nonlinear spatial integration in Off parasol. (right) Summary of discrimination as in left panel across 50 images. Contrast sensitivity was defined as the inverse of the spatial contrast required for a probability of correct discrimination of 0.75.

**Figure 9 - Figure Supplement 2.**
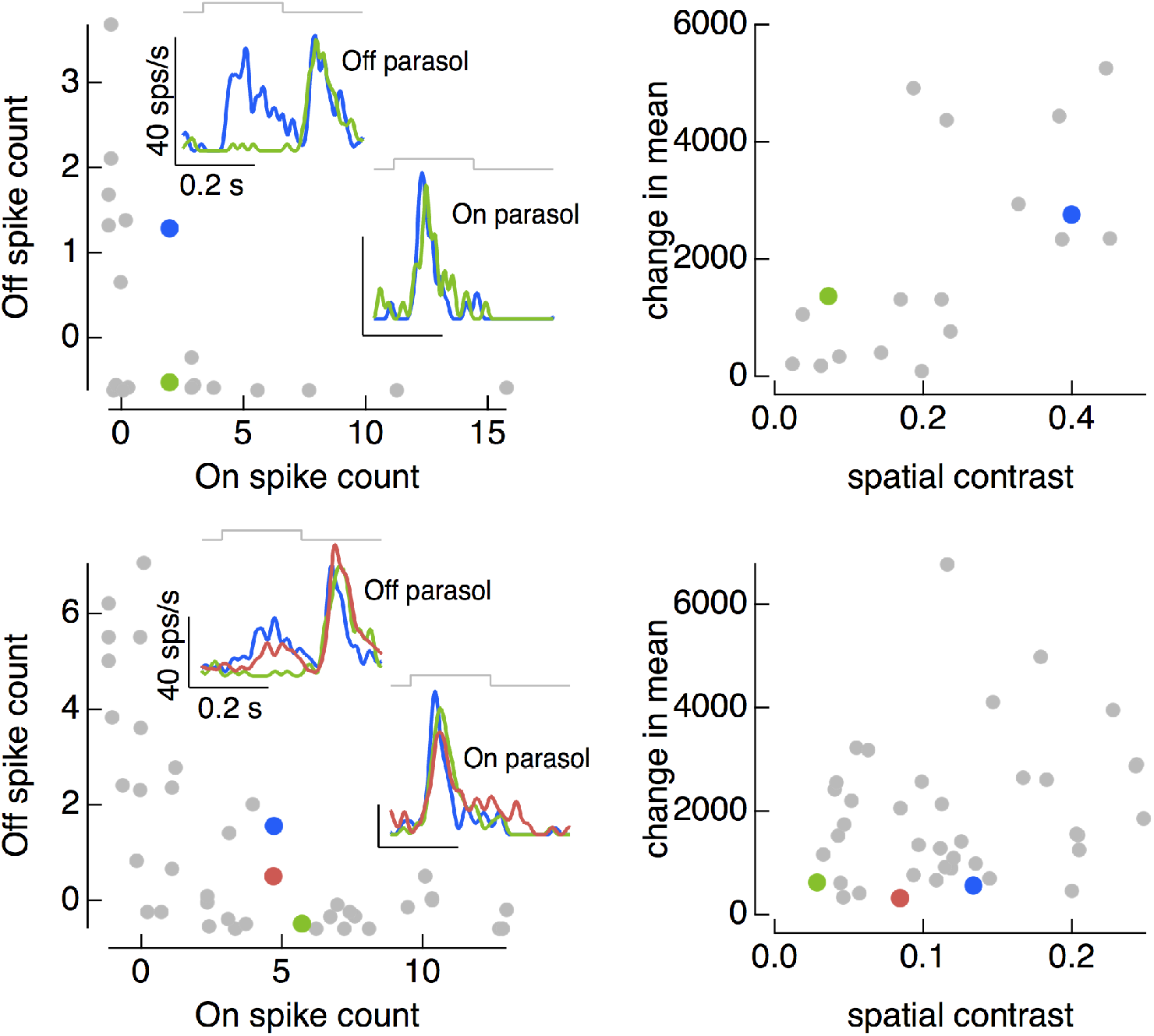
On and Off parasol cell responses to the same collection of image patches for two On/Off pairs. (left) Spike count of responses of Off parasol cell plotted against that of On parasol for a collection of image patches. Insets show highlighted patches. (right) Mean and spatial contrast of each of the patches sampled on the left, with example patches highlighted.

